## Supplementary informations for "Atlasing white matter and grey matter joint contributions to resting-state networks in the human brain"

### Supplementary methods

#### WhiteRest atlas anatomical labelling

We provide a detailed anatomical labelling (white and grey matter) of each resting-state network (RSN) (Supp. Fig. 1 to 30). The RSNs are displayed in 3D, with a “glass brain” effect to show both grey and white matter parts of the RSNs. Multiple views of each RSN are given, with the orientation noted in grey under each view.
In the figures, the grey matter part of the RSN, showed in red, was annotated using the Atlas of Human Brain Connections[1]. Likewise, the white matter part of the RSN showed in green was annotated using the atlas derived from Rojkova et al. (2016)[2]. The labelling was done manually and checked by multiple experts (VN, SF, MTdS, MJ).

#### Dimensionality reduction of clinical scores

As explained in the main manuscript, the neurobehavioral deficit scores used in the study analyses are derived from a principal component analysis (PCA) of neurobehavioral assessment scores (one PCA per deficit). The three neurobehavioral deficits explored where left upper-limb motor control deficit (MotorL deficit), right upper-limb motor control deficit (MotorR), and language deficit. The neurobehavioral assessment scores were associated with each deficit as per the original paper describing the dataset, from Corbetta et al. (2015)[3]:

MotorL deficit:

- Action Research Arm (ARA) test - Left hand grasp
- ARA - Left hand grip
- ARA - Left hand pinch
- Jamar Dynamometer grip strength assessment – Left hand
- 9-Hole Peg test – Left hand
- Left shoulder flexion assessment
- Left wrist extension assessment

MotorR deficit:

- ARA - Right hand grasp
- ARA - Right hand grip
- ARA - Right hand pinch
- Jamar Dynamometer grip strength assessment – Right hand
- 9-Hole Peg test – Right hand
- Right shoulder flexion assessment
- Right wrist extension assessment

---

[1] M. Catani, M. Thiebaut de Schotten (2012). DOI: 10.1093/med/9780199541164.001.0001
[2] K. Rojkova et al. (2016). DOI: 10.1007/s00429-015-1001-3. Atlas available at https://storage.googleapis.com/bcblabweb/open_data.html
[3] M. Corbetta et al. (2015). DOI: 10.1016/j.neuron.2015.02.027

Language deficit:

- Semantic verbal fluency test (SVFT) – Animal name fluency
- Boston Diagnostic Aphasia Examination (BDAE) - Picture naming
- BDAE - Performing listened commands
- BDAE - Nonword repetition
- BDAE - Comprehension of read sentences
- BDAE - Sentence reading
- BDAE - Comprehension of listened word

#### Stroke lesion and RSNs in patients with strong symptoms

To provide an intuitive view of the relationship between lesions, RSN, and symptoms, we selected all the patients who showed a "strong" cognitive deficit and plotted together their lesion and the RSN corresponding to the deficit.

A "strong" deficit for one specific cognitive function was defined as having a (PCA-derived) deficit score in the upper decile of that deficit score (see "*Stroke data analysis*" in the "*Methods*" section of the main manuscript): MotorL deficit ≥ 1; MotorR deficit ≥ 0.97; Language deficit ≥ 0.61. The distribution of these deficit scores and the threshold are displayed in Supp. Fig. 31.

As in the main manuscript, we associated left and right upper-ling motor control deficit with the somato-motor networks for left and right hand, respectively, and the language deficit with the language production network and with the language comprehension network. Each lesion was plotted along the associated RSN (thresholded at z=7), revealing the association between the lesion overlap of the RSN and the corresponding symptoms (Supp. Fig. 32 to 37).

#### Overlap of RSNs and plurality of symptoms

To illustrate the effect of overlapping white matter RSNs on the plurality of symptoms, we selected the patients for whom at least a third (DiscROver score > 33) of both the right hand RSN and the language comprehension RSN were impacted (n = 11). Among them, most (n = 9) showed a "clear" cognitive deficit for both language and right upper-limb motor control ("MotorR deficit"). A "clear" deficit was defined as having a deficit score in the upper quartile of the distribution: Language deficit score ≥ 0.37 and MotorR deficit score ≥ 0.18. These thresholds are displayed on Supp. Fig. 31, and for each of these 9 patients, we displayed the lesion and the two studied network side-by-side in Supp. Fig. 38 and 39.

### Supplementary figures

#### Annotated WhiteRest networks

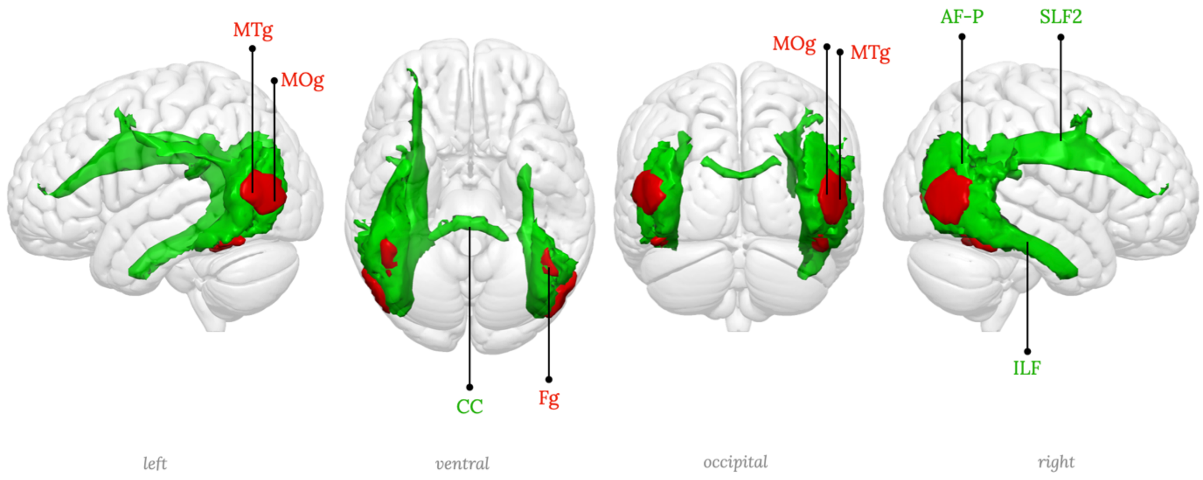

| | | | |
|---|---|---|---|
| Fg | Fusiform gyrus | AF-P | Arcuate fasciculus - posterior segment |
| MTg | Middle temporal gyrus | CC | Corpus callosum |
| MOg | Middle occipital gyrus | ILF | Inferior longitudinal fasciculus |
| | | SLF2 | Second branch of the superior longitudinal fasciculus |

**Supplementary figure 1:** RSN01, Lateral occipital network

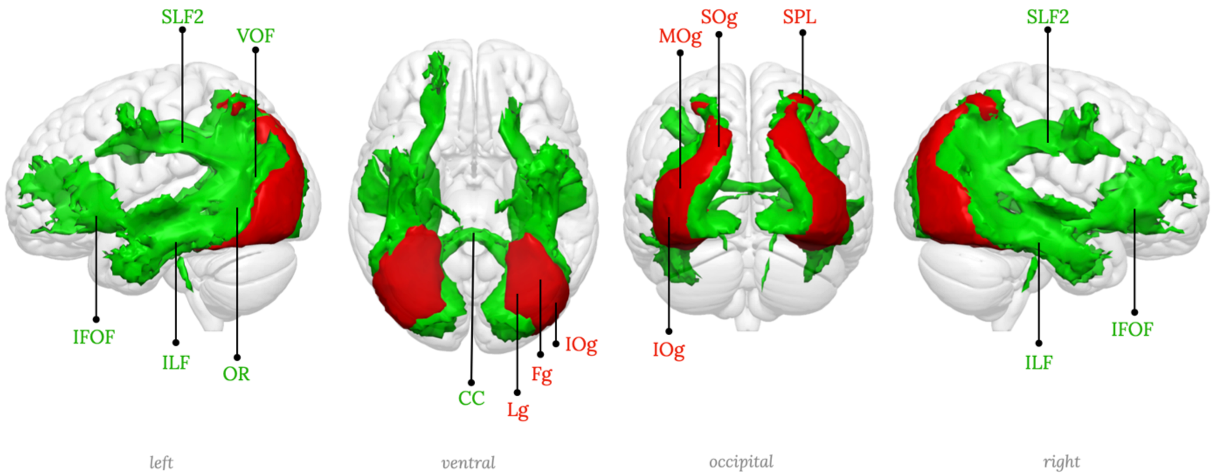

| | | | | | | | |
|---|---|---|---|---|---|---|---|
| Fg | Fusiform gyrus | MOg | Middle occipital gyrus | CC | Corpus callosum | OR | Orbital radiation |
| IOg | Inferior occipital gyrus | SOg | Superior occipital gyrus | IFOF | Inferior fronto-occipital fasciculus | SLF2 | Second branch of the superior longitudinal fasciculus |
| Lg | Lingual gyrus | SPL | Superior parietal lobule | ILF | Inferior longitudinal fasciculus | VOF | Vertical occipital fasciculus |

**Supplementary figure 2:** RSN02, Lateral posterior occipital network

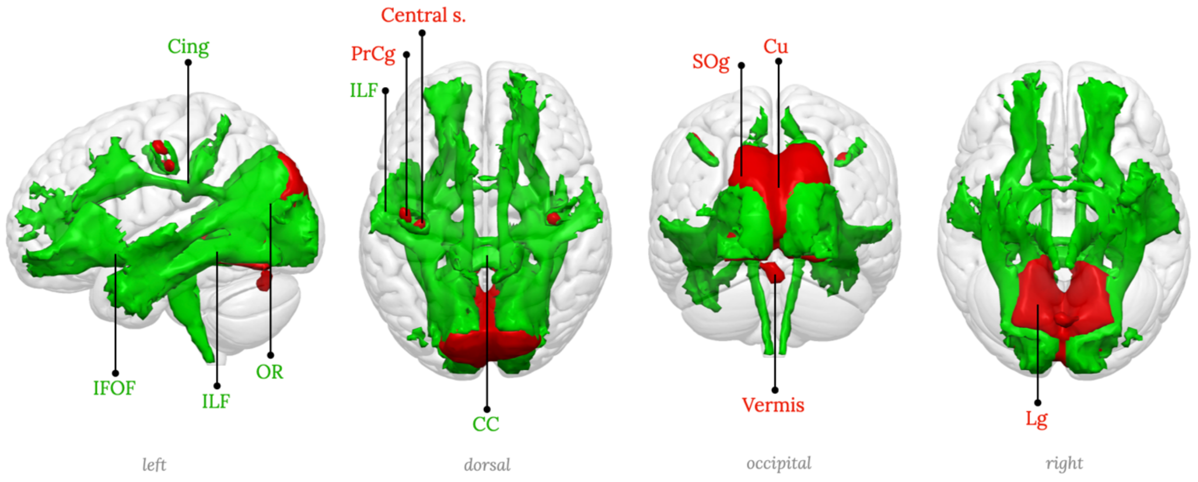

| | | | | | | | |
|---|---|---|---|---|---|---|---|
| Central s. | Central sulcus | PrCg | Precentral gyrus | CC | Corpus callosum | ILF | Inferior longitudinal fasciculus |
| Cu | Cuneus | SOg | Superior occipital gyrus | Cing | Cingulum | OR | Orbital radiation |
| Lg | Lingual gyrus | | | IFOF | Inferior fronto-occipital fasciculus | | |

**Supplementary figure 3:** RSN03, Medial occipital network

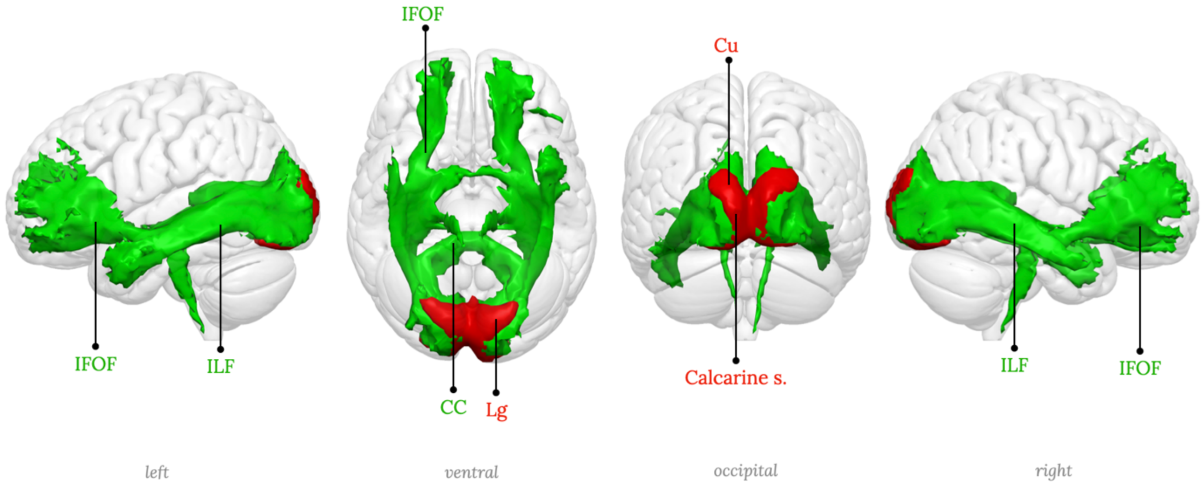

| | | | |
|---|---|---|---|
| Calcarine s. | Calcarine sulcus | CC | Corpus callosum |
| Cu | Cuneus | IFOF | Inferior fronto-occipital fasciculus |
| Lg | Lingual gyrus | ILF | Inferior longitudinal fasciculus |

**Supplementary figure 4:** RSN04, Medial posterior occipital network, primary visual network

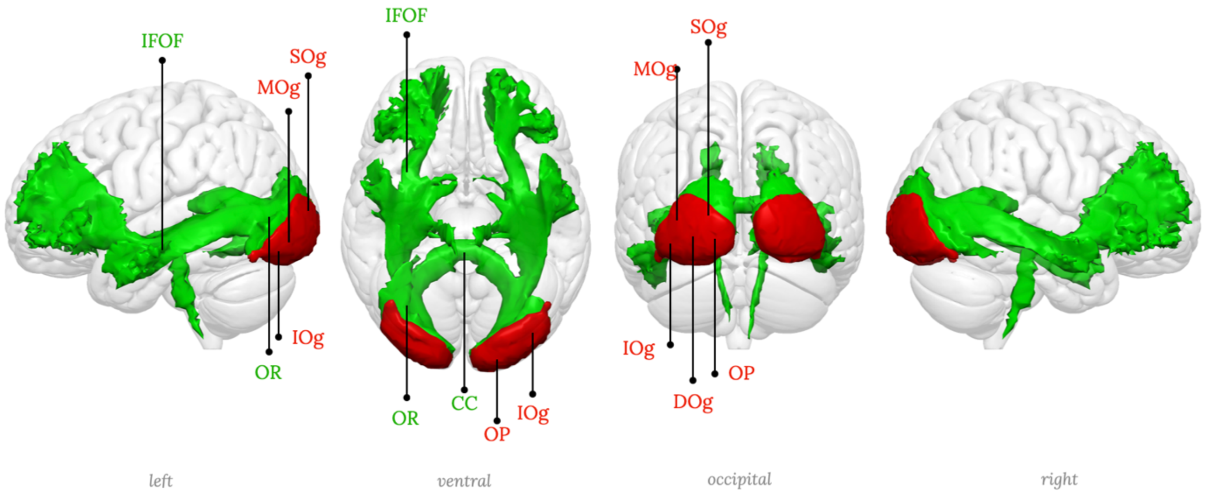

| | | | | | |
|---|---|---|---|---|---|
| DOg | Descending occipital gyrus | OP | Occipital pole | CC | Corpus callosum |
| IOg | Inferior occipital gyrus | SOg | Superior occipital gyrus | IFOF | Inferior fronto-occipital fasciculus |
| MOg | Middle occipital gyrus | | | OR | Orbital radiation |

**Supplementary figure 5:** RSN05, Posterior occipital network.

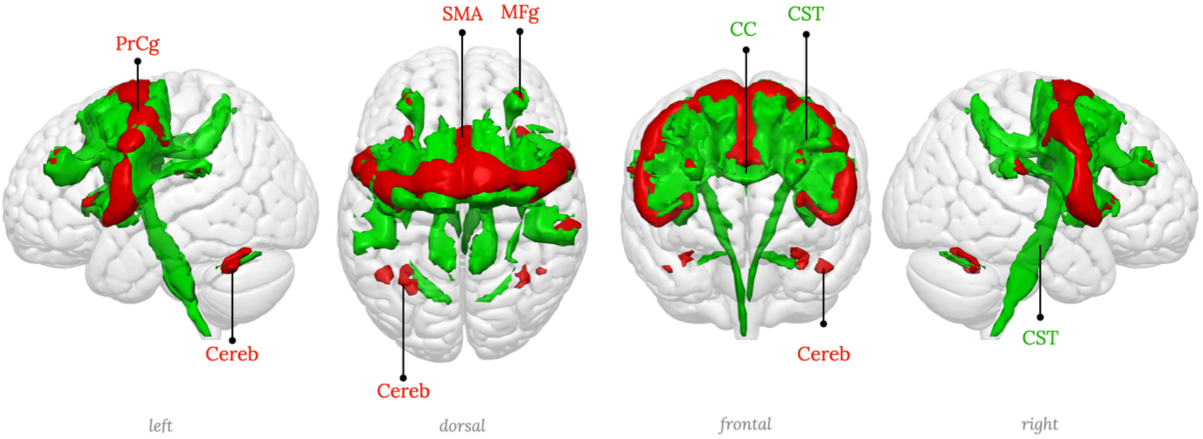

| | | | | | |
|---|---|---|---|---|---|
| Cereb | Cerebellum | SMA | Supplementary motor area | CC | Corpus callosum |
| MFg | Middle frontal gyrus | | | CST | Corticospinal tract |
| PrCg | Precentral gyrus | | | | |

**Supplementary figure 6:** RSN06, Precentral network, Motor network.

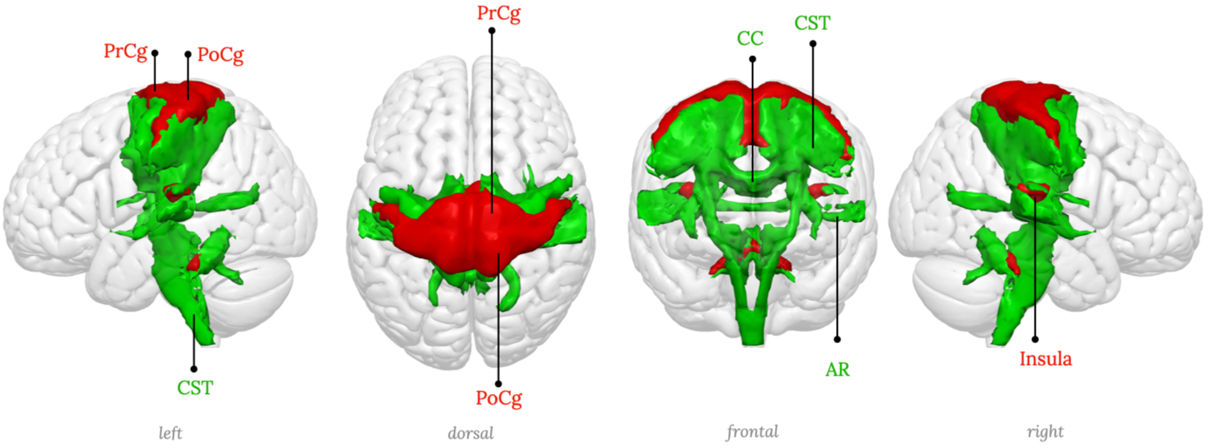

| | | | |
|---|---|---|---|
| PoCg | Postcentral gyrus | AR | Acoustic radiation |
| PrCg | Precentral gyrus | CC | Corpus callosum |
| | | CST | Corticospinal tract |

**Supplementary figure 7:** RSN07, Middle central network, left hemisphere component (somato-motor, right hand portion)

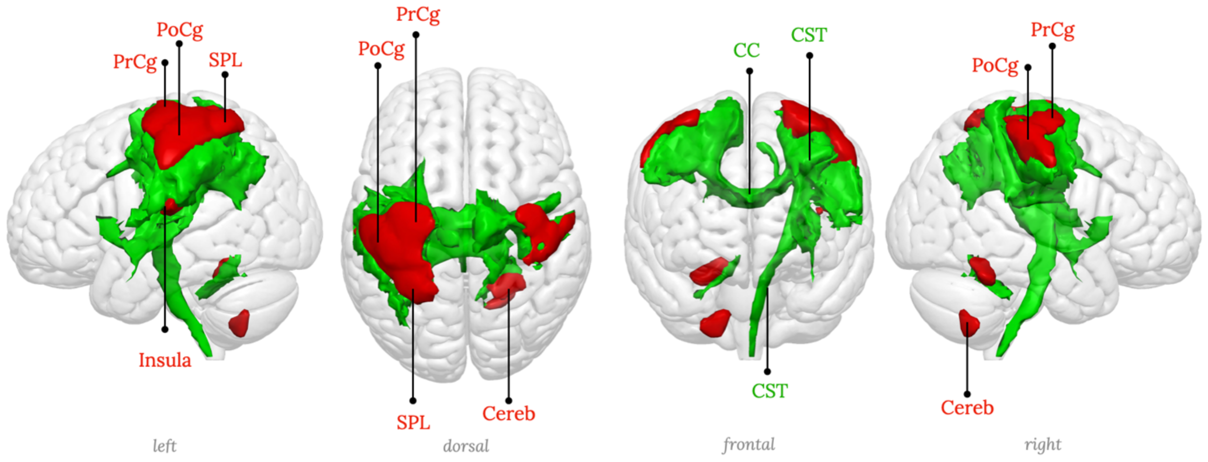

| | | | |
|---|---|---|---|
| Cereb | Cerebellum | CC | Corpus callosum |
| PoCg | Postcentral gyrus | CST | Corticospinal tract |
| PrCg | Precentral gyrus | | |
| SPL | Superior parietal lobule | | |

**Supplementary figure 8:** RSN08, Middle central network, left hemisphere component (somato-motor, right hand portion)

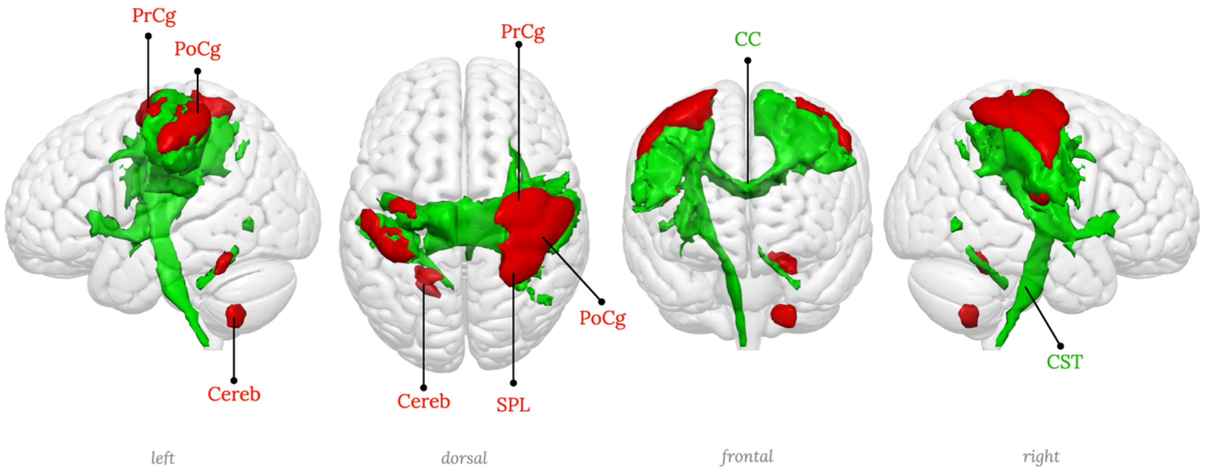

| | | | |
|---|---|---|---|
| Cereb | Cerebellum | CC | Corpus callosum |
| PoCg | Postcentral gyrus | CST | Corticospinal tract |
| PrCg | Precentral gyrus | | |
| SPL | Superior parietal lobule | | |

**Supplementary figure 9:** RSN09, Middle central network, right hemisphere component (somato-motor, left hand portion)

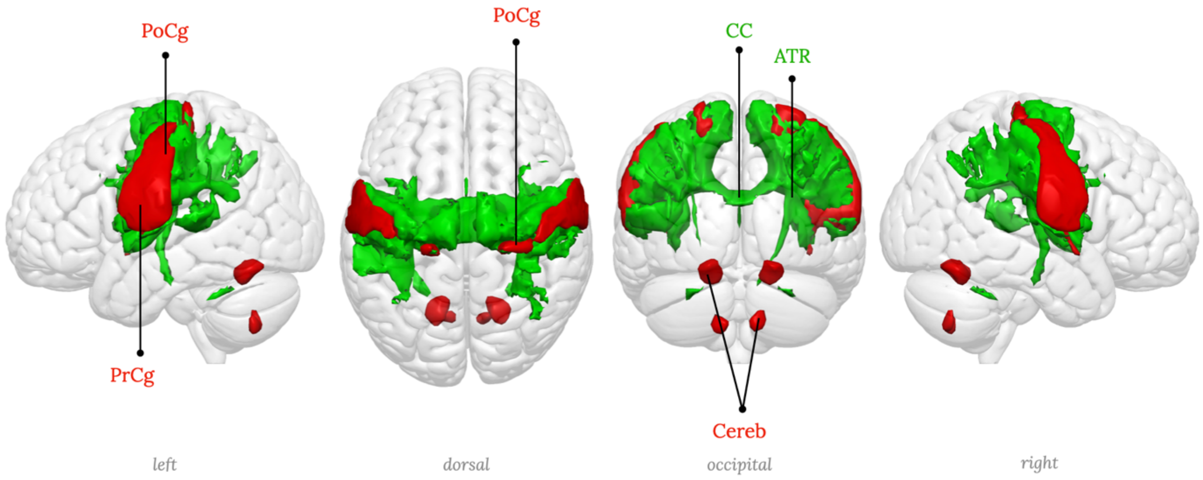

| | | | |
|---|---|---|---|
| Cereb | Cerebellum | ATR | Anterior thalamic radiation |
| PoCg | Postcentral gyrus | CC | Corpus callosum |
| PrCg | Precentral gyrus | | |

**Supplementary figure 10:** RSN10, Inferior central network (somato-motor, head portion)

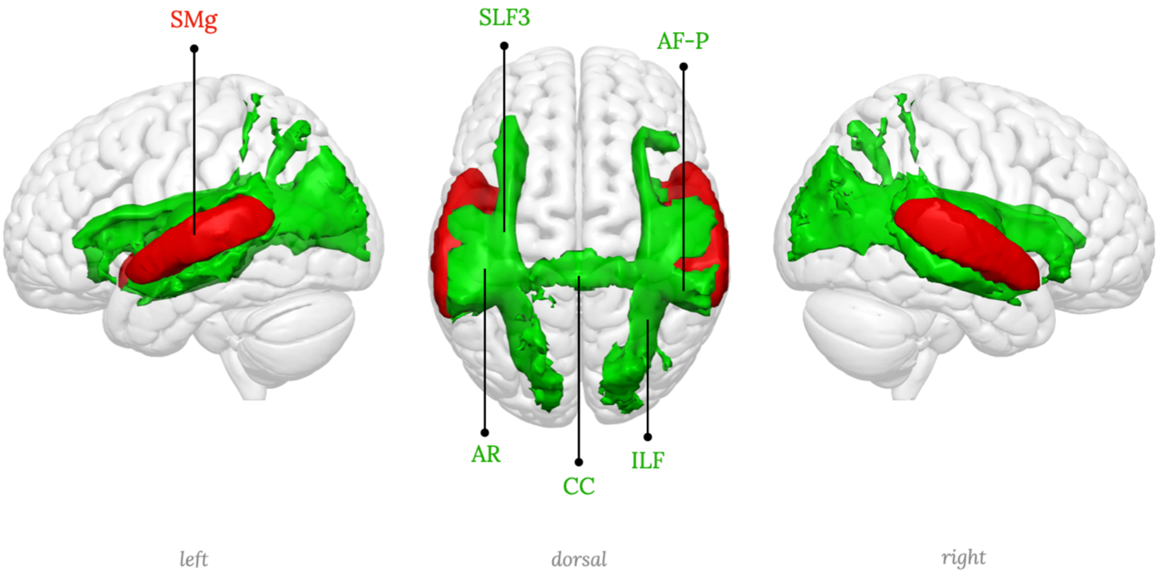

| | |
|---|---|
| SMg | Supramarginal gyrus |

| | | | |
|---|---|---|---|
| AF-P | Arcuate fasciculus - posterior segment | ILF | Inferior longitudinal fasciculus |
| AR | Acoustic radiation | SLF3 | Third branch of the superior longitudinal fasciculus |
| CC | Corpus callosum | | |

**Supplementary figure 11:** RSN11, Superior temporal network, Primary auditory network.

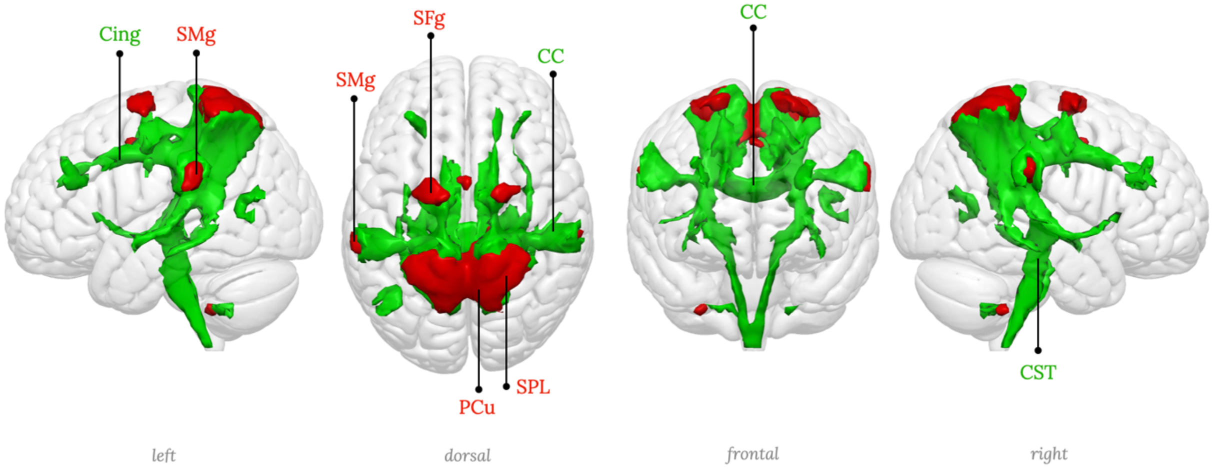

| | | | |
|---|---|---|---|
| PCu | Precuneus | CC | Corpus callosum |
| SFg | Superior frontal gyrus | Cing | Cingulum |
| SMg | Supramarginal gyrus | CST | Corticospinal tract |
| SPL | Superior parietal lobule | | |

**Supplementary figure 12:** RSN12, Anterior superior parietal network

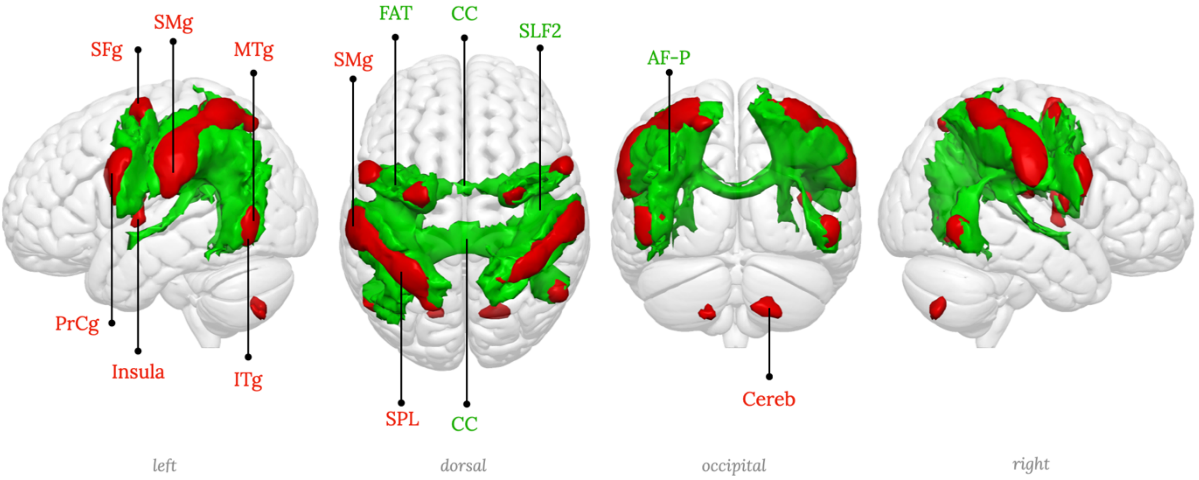

| | | | | | |
|---|---|---|---|---|---|
| Cereb | Cerebellum | SFg | Superior frontal gyrus | AF-P | Arcuate fasciculus - posterior segment |
| ITg | Inferior temporal gyrus | SMg | Supramarginal gyrus | CC | Corpus callosum |
| MTg | Middle temporal gyrus | SPL | Superior parietal lobule | FAT | Frontal aslant tract |
| PrCg | Precentral gyrus | | | SLF2 | Second branch of the superior longitudinal fasciculus |

**Supplementary figure 13:** RSN13, Superior parietal network, Dorsal attention network

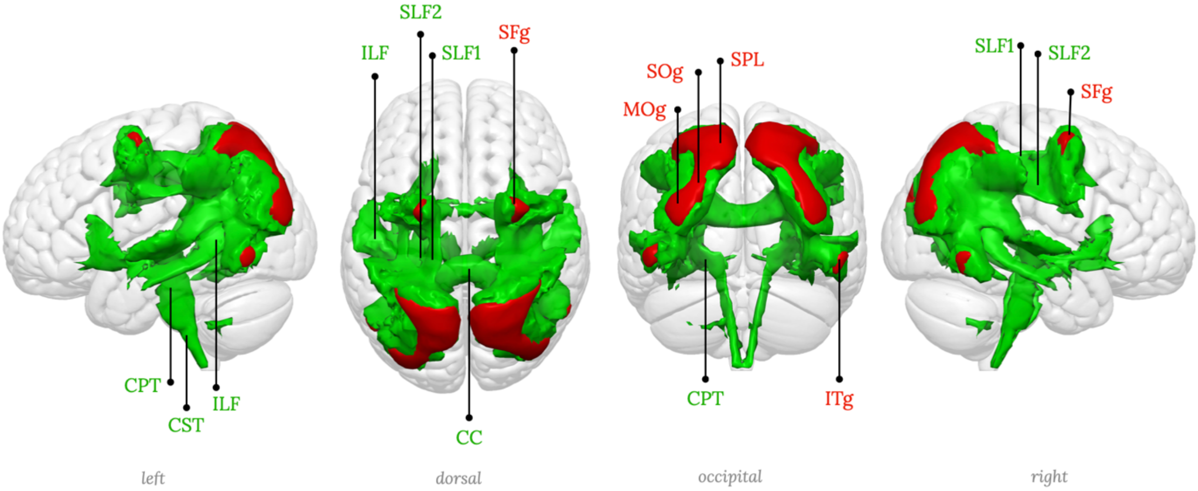

| | | | | | | | |
|---|---|---|---|---|---|---|---|
| ITg | Inferior temporal gyrus | SOg | Superior occipital gyrus | CC | Corpus callosum | ILF | Inferior longitudinal fasciculus |
| MOg | Middle occipital gyrus | SPL | Superior parietal lobule | CPT | Corticpontine tract | SLF1 | First branch of the superior longitudinal fasciculus |
| SFg | Superior frontal gyrus | | | CST | Cortico Spinal tract | SLF2 | Second branch of the superior longitudinal fasciculus |

**Supplementary figure 14:** RSN14, Posterior superior parietal network

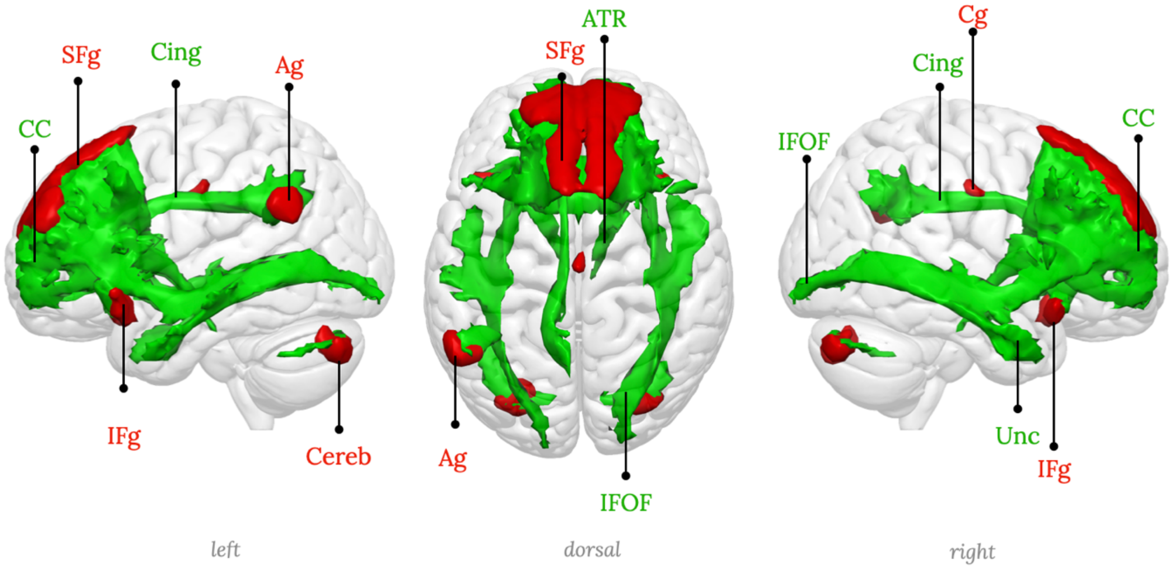

| | | | | | | | |
|---|---|---|---|---|---|---|---|
| Ag | Angular gyrus | IFg | Inferior frontal gyrus | ATR | Anterior thalamic radiation | IFOF | Inferior fronto-occipital fasciculus |
| Cereb | Cerebellum | SFg | Superior frontal gyrus | CC | Corpus callosum | Unc | Uncinate fasciculus |
| Cg | Cingulate gyrus | | | Cing | Cingulum | | |

**Supplementary figure 15:** RSN15, Medial frontal network

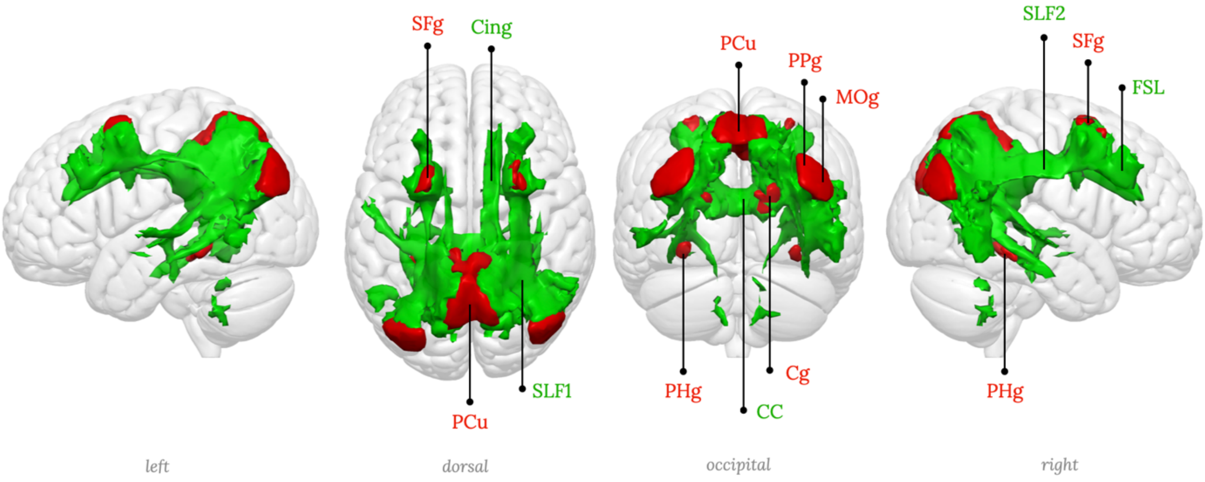

| | | | | | |
|---|---|---|---|---|---|
| Cg | Cingulate gyrus | PHg | Parahippocampal gyrus | CC | Corpus callosum |
| MOg | Middle occipital gyrus | PPg | Posterior parietal gyrus | Cing | Cingulum |
| PCu | Precuneus | SFg | Superior frontal gyrus | FSL | Frontal superior longitudinal tract |
| | | | | SLF2 | Second branch of the superior longitudinal fasciculus |

**Supplementary figure 16:** RSN16, Posterior parietal-precuneal network, Posterior default mode network

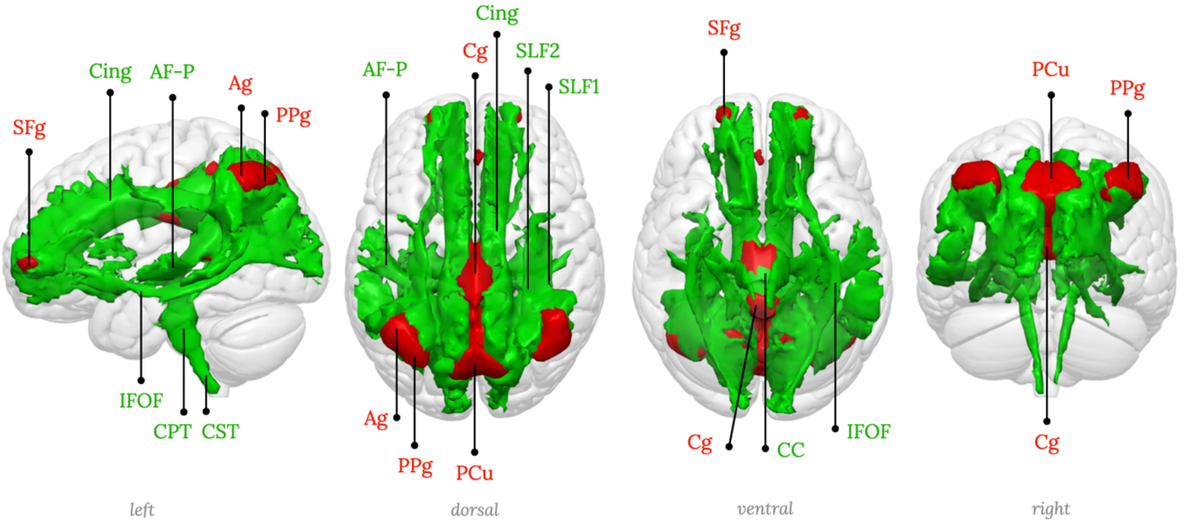

| | | | | | | | |
|---|---|---|---|---|---|---|---|
| Ag | Angular gyrus | PPg | Posterior parietal gyrus | AF-P | Arcuate fasciculus - posterior segment | IFOF | Inferior fronto-occipital fasciculus |
| Cg | Cingulate gyrus | SFg | Superior frontal gyrus | CC | Corpus callosum | SLF1 | First branch of the superior longitudinal fasciculus |
| PCu | Precuneus | | | Cing | Cingulum | SLF2 | Second branch of the superior longitudinal fasciculus |
| | | | | CPT | Corticpontine tract | | |

**Supplementary figure 17:** RSN17, Posterior cingulate-precuneal network, Dorsal default mode network

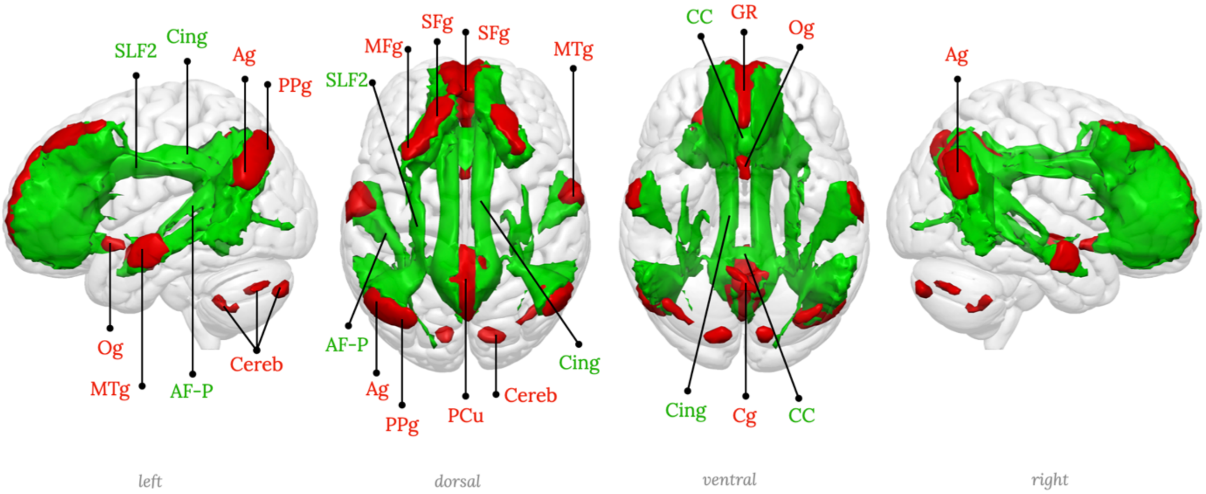

| | | | | | |
|---|---|---|---|---|---|
| Ag | Angular gyrus | MFg | Middle frontal gyrus | PPg | Posterior parietal gyrus |
| Cereb | Cerebellum | MTg | Middle temporal gyrus | SFg | Superior frontal gyrus |
| Cg | Cingulate gyrus | Og | Olfactory gyrus | | |
| GR | Gyrus rectus | PCu | Precuneus | | |

| | |
|---|---|
| AF-P | Arcuate fasciculus - posterior segment |
| CC | Corpus callosum |
| Cing | Cingulum |
| SLF2 | Second branch of the superior longitudinal fasciculus |

**Supplementary figure 18:** RSN18, Precuneus network, Default mode network proper

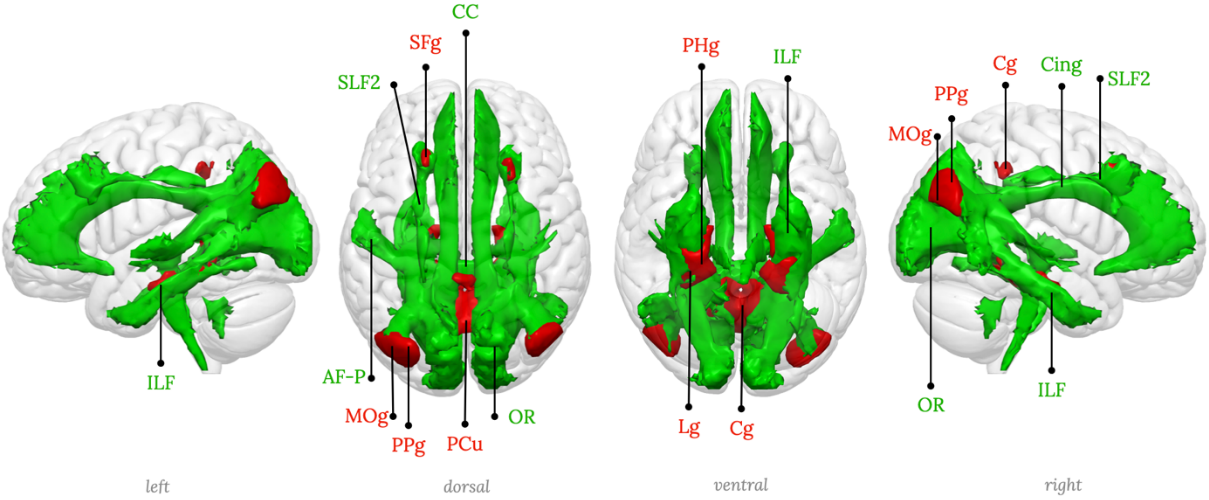

| | | | |
|---|---|---|---|
| Cg | Cingulate gyrus | PHg | Parahippocampal gyrus |
| Lg | Lingual gyrus | PPg | Posterior parietal gyrus |
| MOg | Middle occipital gyrus | SFg | Superior frontal gyrus |
| PCu | Precuneus | | |

| | | | |
|---|---|---|---|
| AF-P | Arcuate fasciculus - posterior segment | ILF | Inferior longitudinal fasciculus |
| CC | Corpus callosum | OR | Orbital radiation |
| Cing | Cingulum | SLF2 | Second branch of the superior longitudinal fasciculus |

**Supplementary figure 19:** RSN19, Parahippocampal-precuneal network, Parahippocampal default mode network

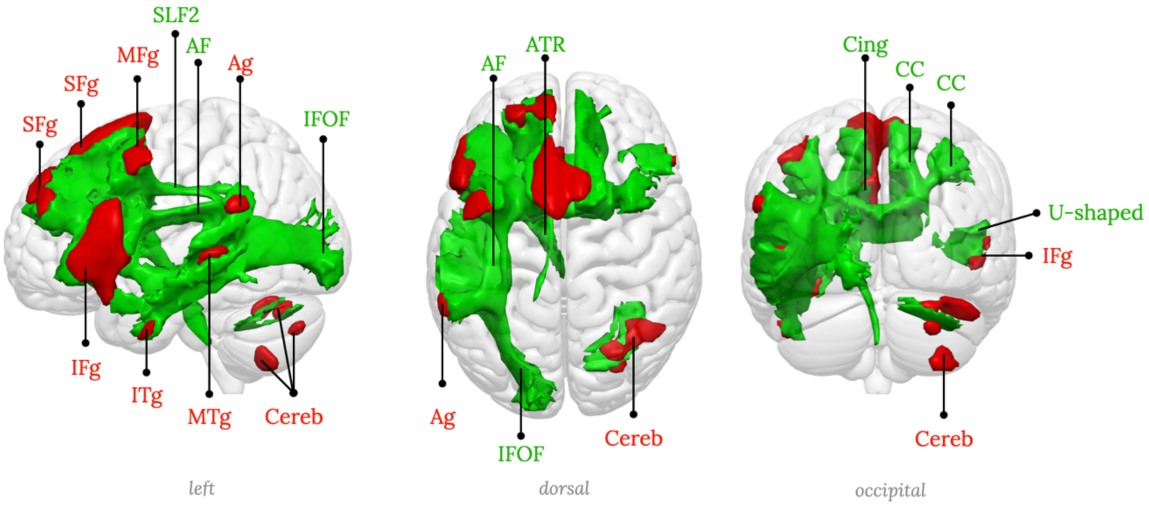

| | | | |
|---|---|---|---|
| Ag | Angular gyrus | MFg | Middle frontal gyrus |
| Cereb | Cerebellum | MTg | Middle temporal gyrus |
| IFg | Inferior frontal gyrus | SFg | Superior frontal gyrus |
| ITg | Inferior temporal gyrus | | |

| | | | |
|---|---|---|---|
| AF | Arcuate fasciculus | Cing | Cingulum |
| ATR | Anterior thalamic radiation | IFOF | Inferior fronto-occipital fasciculus |
| CC | Corpus callosum | SLF2 | Second branch of the superior longitudinal fasciculus |

**Supplementary figure 20:** RSN20, Left inferior frontal network, Language production network

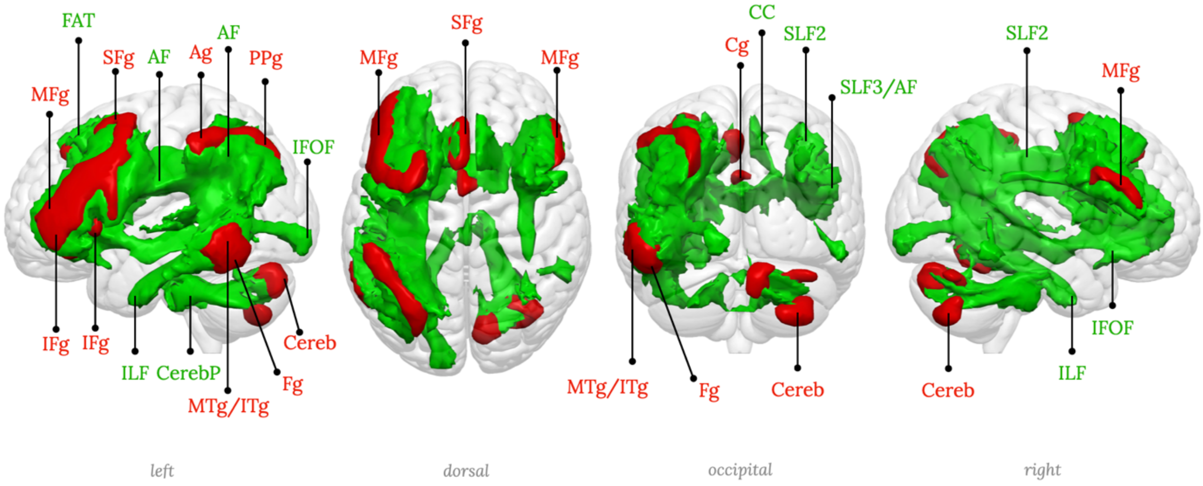

| | | | | | |
|---|---|---|---|---|---|
| Ag | Angular gyrus | IFg | Inferior frontal gyrus | PPg | Posterior parietal gyrus |
| Cereb | Cerebellum | ITg | Inferior temporal gyrus | SMg | Supramarginal gyrus |
| Cg | Cingulate gyrus | MFg | Middle frontal gyrus | | |
| Fg | Fusiform gyrus | MTg | Middle temporal gyrus | | |

| | | | |
|---|---|---|---|
| AF | Arcuate fasciculus | FAT | Frontal aslant tract |
| CC | Corpus callosum | ILF | Inferior longitudinal fasciculus |
| CerebP | Pontocerebellar tract | SLF2 | Second branch of the superior longitudinal fasciculus |
| | | SLF3 | Third branch of the superior longitudinal fasciculus |

**Supplementary figure 21:** RSN21, Fronto-parieto-temporal network 1, left hemisphere component

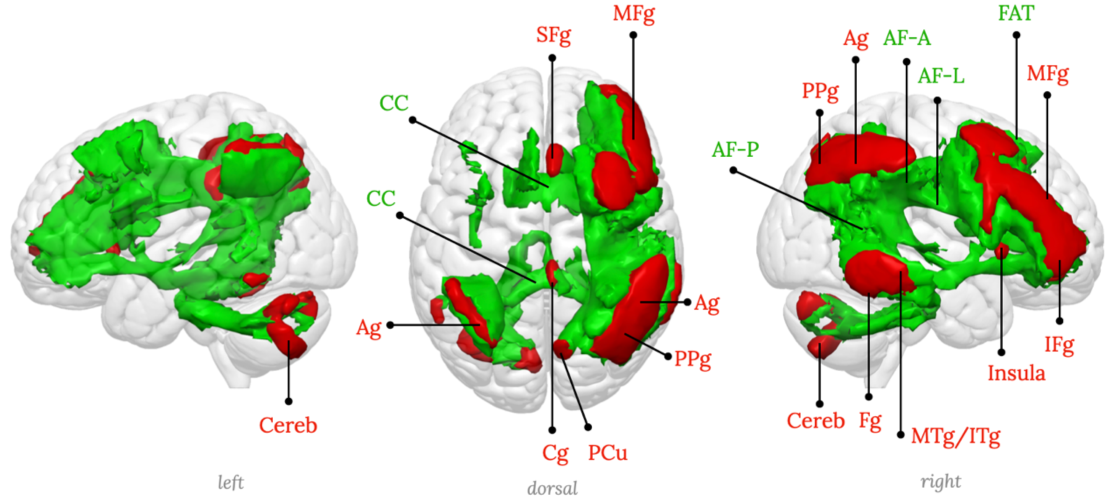

| | | | | | |
|---|---|---|---|---|---|
| Ag | Angular gyrus | IFg | Inferior frontal gyrus | PCu | Precuneus |
| Cereb | Cerebellum | ITg | Inferior temporal gyrus | PPg | Posterior parietal gyrus |
| Cg | Cingulate gyrus | MFg | Middle frontal gyrus | SFg | Superior frontal gyrus |
| Fg | Fusiform gyrus | MTg | Middle temporal gyrus | | |

| | | | |
|---|---|---|---|
| AF-A | Arcuate fasciculus - anterior segment | CC | Corpus callosum |
| AF-L | Arcuate fasciculus - long segment | FAT | Frontal aslant tract |
| AF-P | Arcuate fasciculus - posterior segment | | |

**Supplementary figure 22:** RSN22, Fronto-parieto-temporal network 1, right hemisphere component

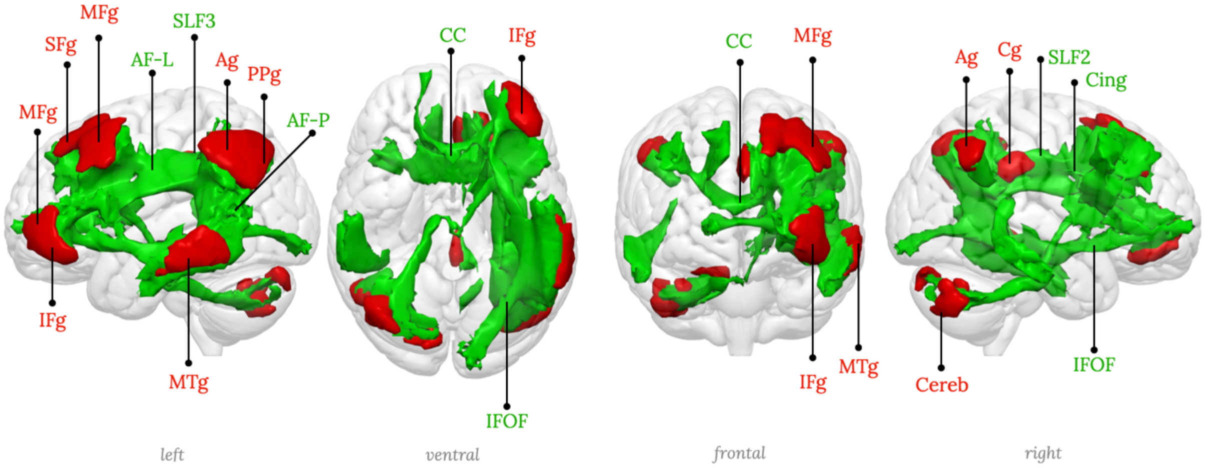

| | | | |
|---|---|---|---|
| Ag | Angular gyrus | MFg | Middle frontal gyrus |
| Cereb | Cerebellum | MTg | Middle temporal gyrus |
| Cg | Cingulate gyrus | PPg | Posterior parietal gyrus |
| IFg | Inferior frontal gyrus | SFg | Superior frontal gyrus |

| | | | |
|---|---|---|---|
| AF-L | Arcuate fasciculus - long segment | IFOF | Inferior fronto-occipital fasciculus |
| AF-P | Arcuate fasciculus - posterior segment | SLF2 | Second branch of the superior longitudinal fasciculus |
| CC | Corpus callosum | SLF3 | Third branch of the superior longitudinal fasciculus |
| Cing | Cingulum | | |

**Supplementary figure 23:** RSN23, Fronto-parieto-temporal network 2, left hemisphere component

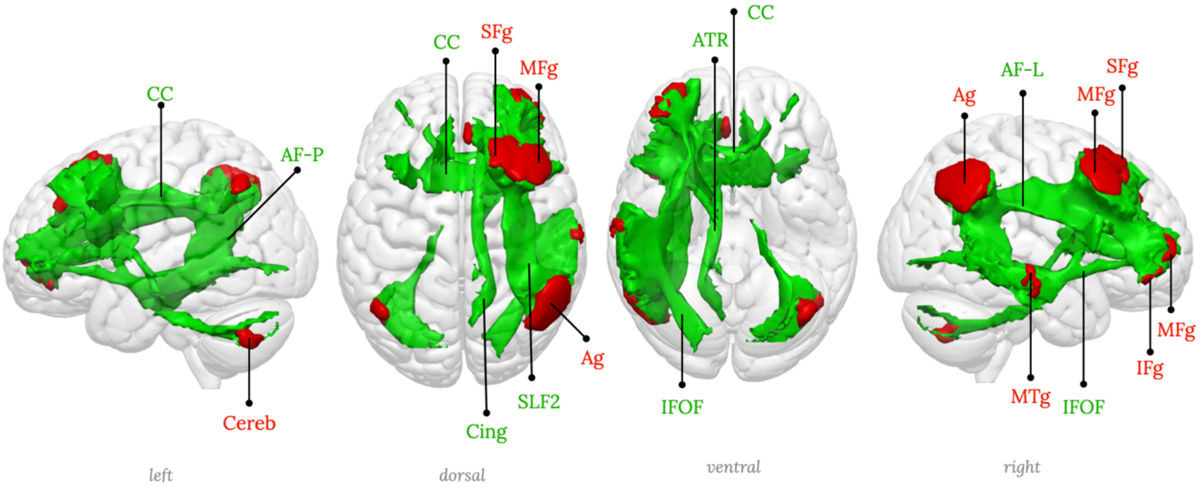

| | | | |
|---|---|---|---|
| Ag | Angular gyrus | MFg | Middle frontal gyrus |
| Cereb | Cerebellum | MTg | Middle temporal gyrus |
| IFg | Inferior frontal gyrus | SFg | Superior frontal gyrus |

| | | | |
|---|---|---|---|
| AF-L | Arcuate fasciculus - long segment | Cing | Cingulum |
| AF-P | Arcuate fasciculus - posterior segment | IFOF | Inferior fronto-occipital fasciculus |
| ATR | Anterior thalamic radiation | SLF2 | Second branch of the superior longitudinal fasciculus |
| CC | Corpus callosum | | |

**Supplementary figure 24:** RSN24, Fronto-parieto-temporal network 2, right hemisphere component

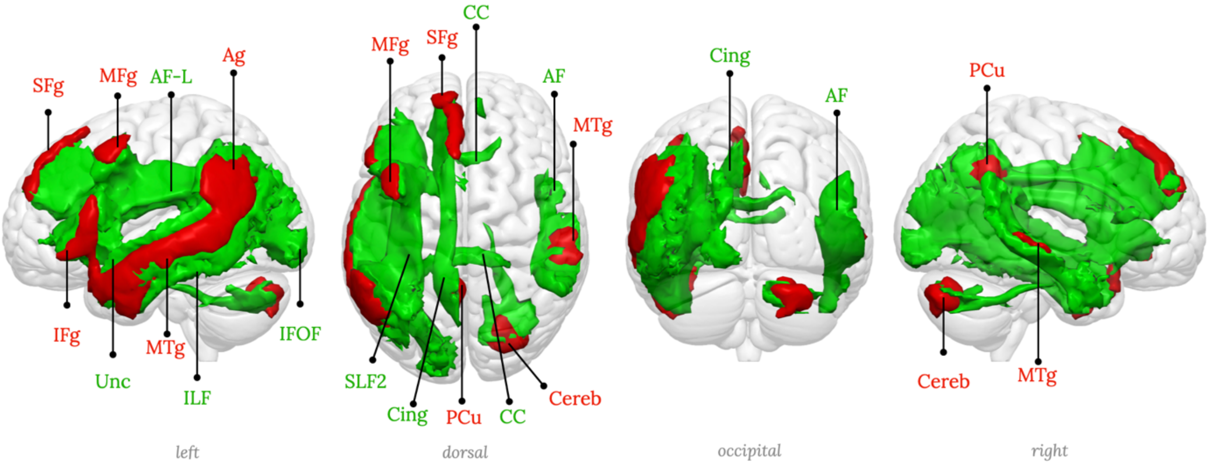

| | | | |
|---|---|---|---|
| Ag | Angular gyrus | MTg | Middle temporal gyrus |
| Cereb | Cerebellum | PCu | Precuneus |
| IFg | Inferior frontal gyrus | SFg | Superior frontal gyrus |
| MFg | Middle frontal gyrus | | |

| | | | |
|---|---|---|---|
| AF | Arcuate fasciculus | IFOF | Inferior fronto-occipital fasciculus |
| AF-L | Arcuate fasciculus - long segment | ILF | Inferior longitudinal fasciculus |
| CC | Corpus callosum | SLF2 | Second branch of the superior longitudinal fasciculus |
| Cing | Cingulum | Unc | Uncinate fasciculus |

**Supplementary figure 25:** RSN25, Fronto-parieto-temporal network 3, left hemisphere component, Language comprehension network

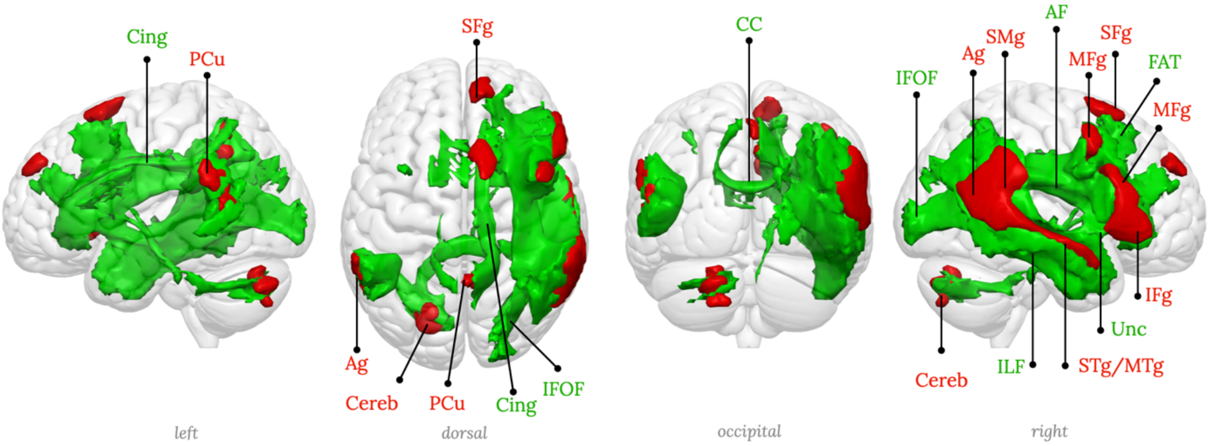

| | | | | | |
|---|---|---|---|---|---|
| Ag | Angular gyrus | MFg | Middle frontal gyrus | SFg | Superior frontal gyrus |
| Cereb | Cerebellum | MTg | Middle temporal gyrus | SMg | Supramarginal gyrus |
| IFg | Inferior frontal gyrus | PCu | Precuneus | STg | Superior temporal gyrus |

| | | | |
|---|---|---|---|
| AF | Arcuate fasciculus | IFOF | Inferior fronto-occipital fasciculus |
| CC | Corpus callosum | ILF | Inferior longitudinal fasciculus |
| Cing | Cingulum | Unc | Uncinate fasciculus |
| FAT | Frontal aslant tract | | |

**Supplementary figure 26:** RSN26, Fronto-parieto-temporal network 3, right hemisphere component

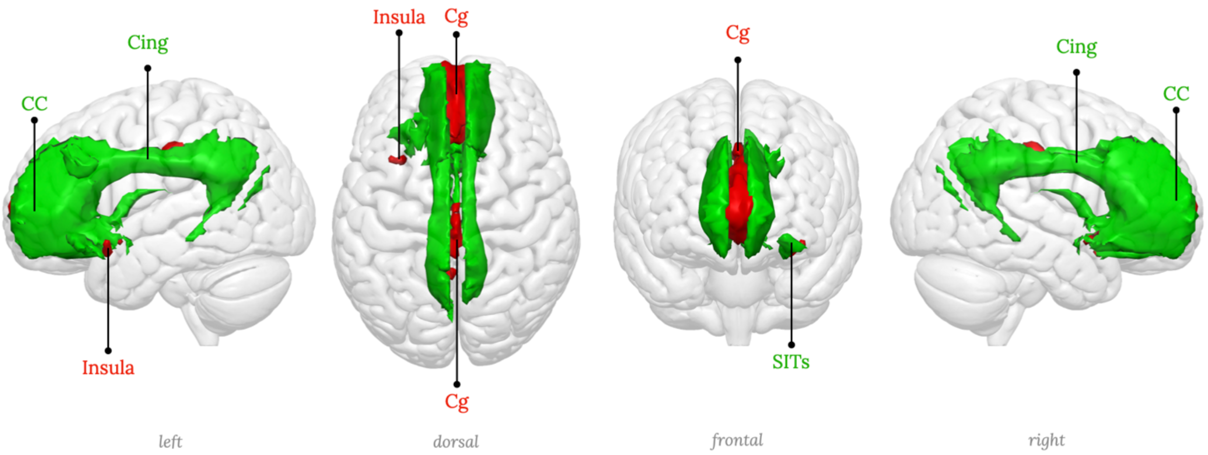

| | | | |
|---|---|---|---|
| | | CC | Corpus callosum |
| Cg | Cingulate gyrus | Cing | Cingulum |
| | | SITs | Small insular tracts |

**Supplementary figure 27:** RSN27, Anterior cingulate network

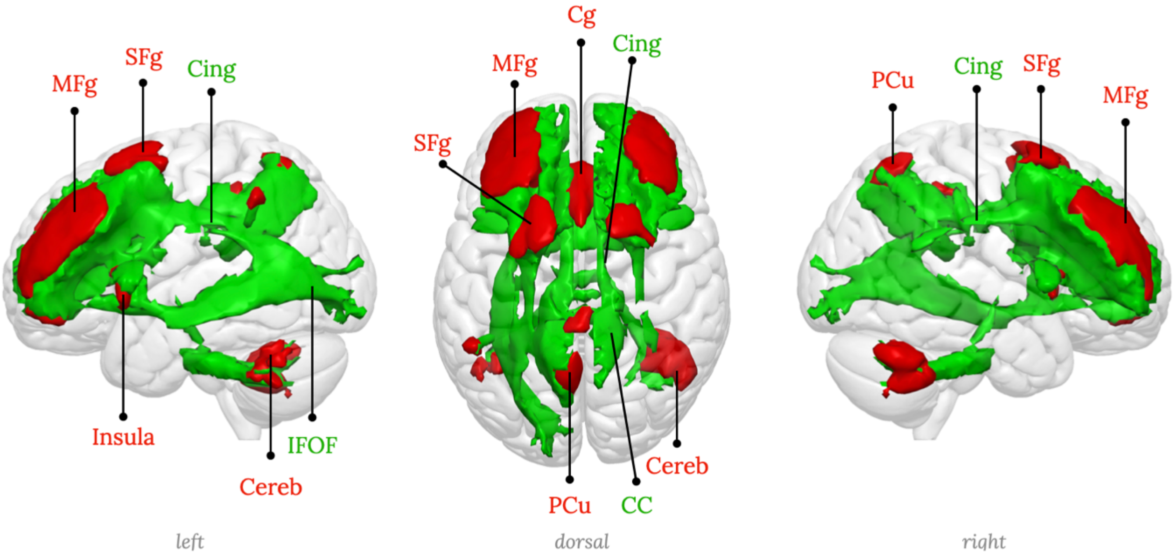

| | | | | | |
|---|---|---|---|---|---|
| Cereb | Cerebellum | PCu | Precuneus | CC | Corpus callosum |
| Cg | Cingulate gyrus | SFg | Superior frontal gyrus | Cing | Cingulum |
| MFg | Middle frontal gyrus | | | IFOF | Inferior fronto-occipital fasciculus |

**Supplementary figure 28:** RSN28, Dorso-lateral prefrontal cortex network

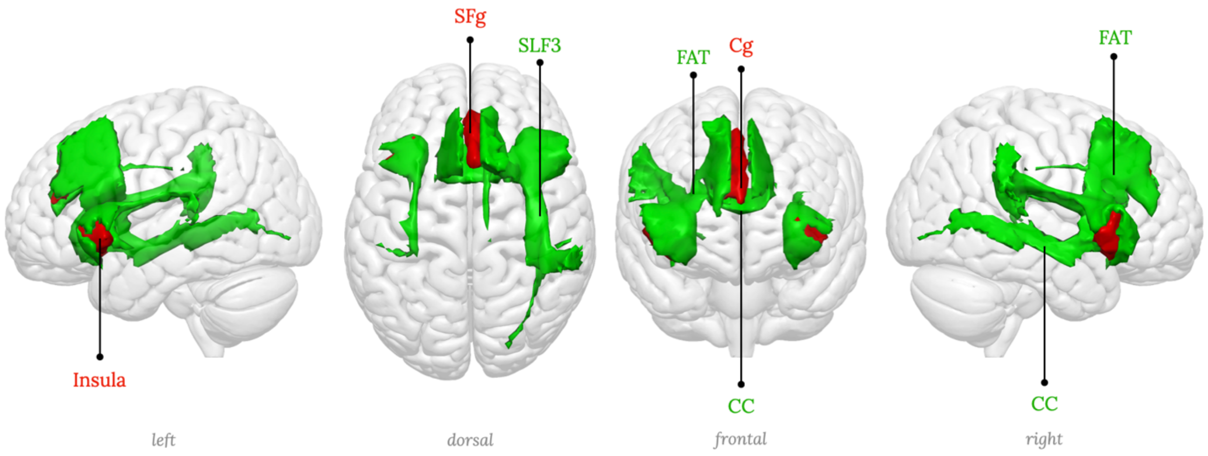

| | | | |
|---|---|---|---|
| Cg | Cingulate gyrus | CC | Corpus callosum |
| SFg | Superior frontal gyrus | FAT | Frontal aslant tract |
| | | SLF3 | Third branch of the superior longitudinal fasciculus |

**Supplementary figure 29:** RSN29, Insula network, Salience network

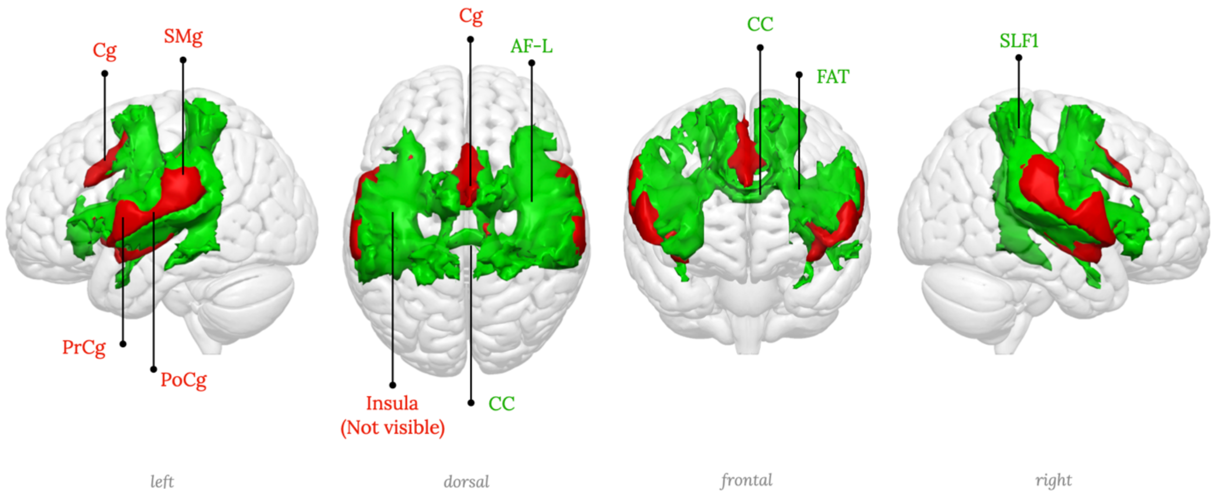

| | | | |
|---|---|---|---|
| Cg | Cingulate gyrus | AF-L | Arcuate fasciculus - long segment |
| PoCg | Postcentral gyrus | CC | Corpus callosum |
| PrCg | Precentral gyrus | FAT | Frontal aslant tract |
| SMg | Supramarginal gyrus | SLF1 | First branch of the superior longitudinal fasciculus |

**Supplementary figure 30:** RSN30, Cingulo-opercular network

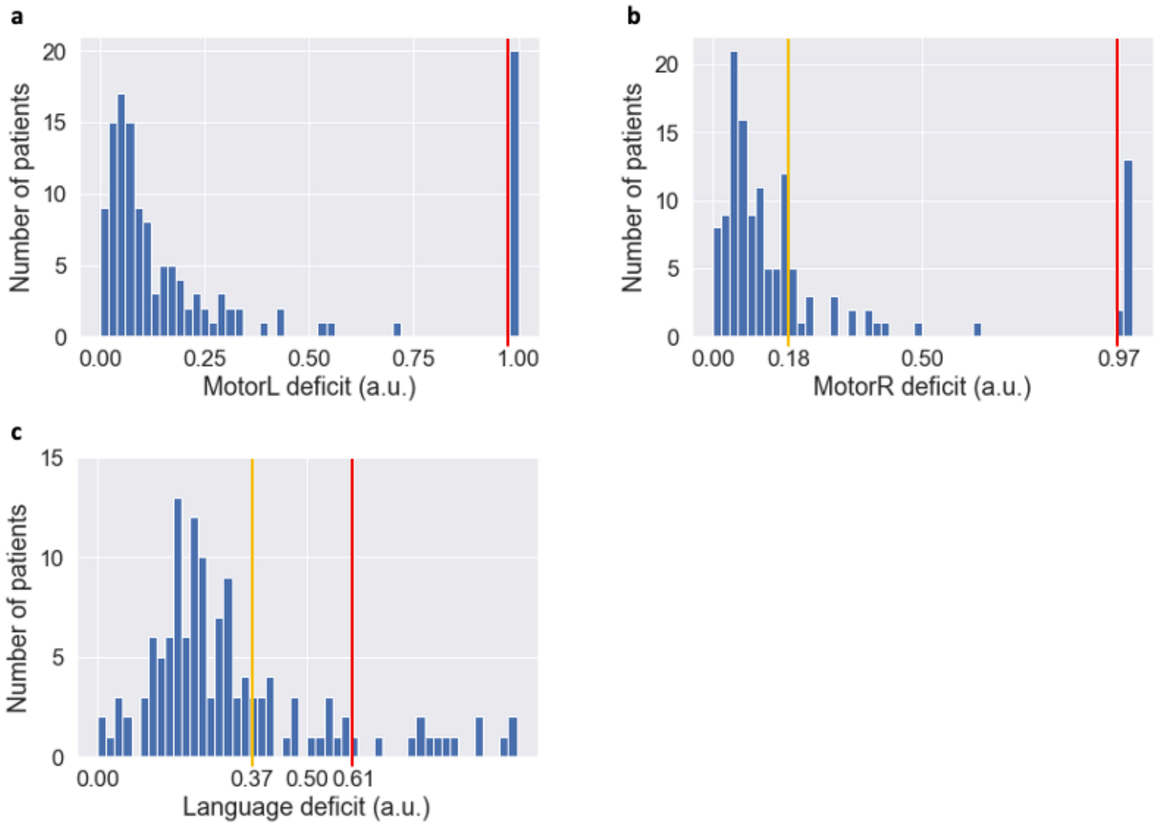

**Supplementary figure 31**: Distribution of cognitive deficit PC1 scores. **a**: Scores for left upper-limb motor control deficit. **b**: Scores for right upper-limb motor control deficit. **c**: Scores for language deficit. The orange bar indicates the upper quartile (i.e., "clear" deficit), and the red bar indicates the upper decile (i.e., "strong" deficit). a.u.: arbitrary unit.

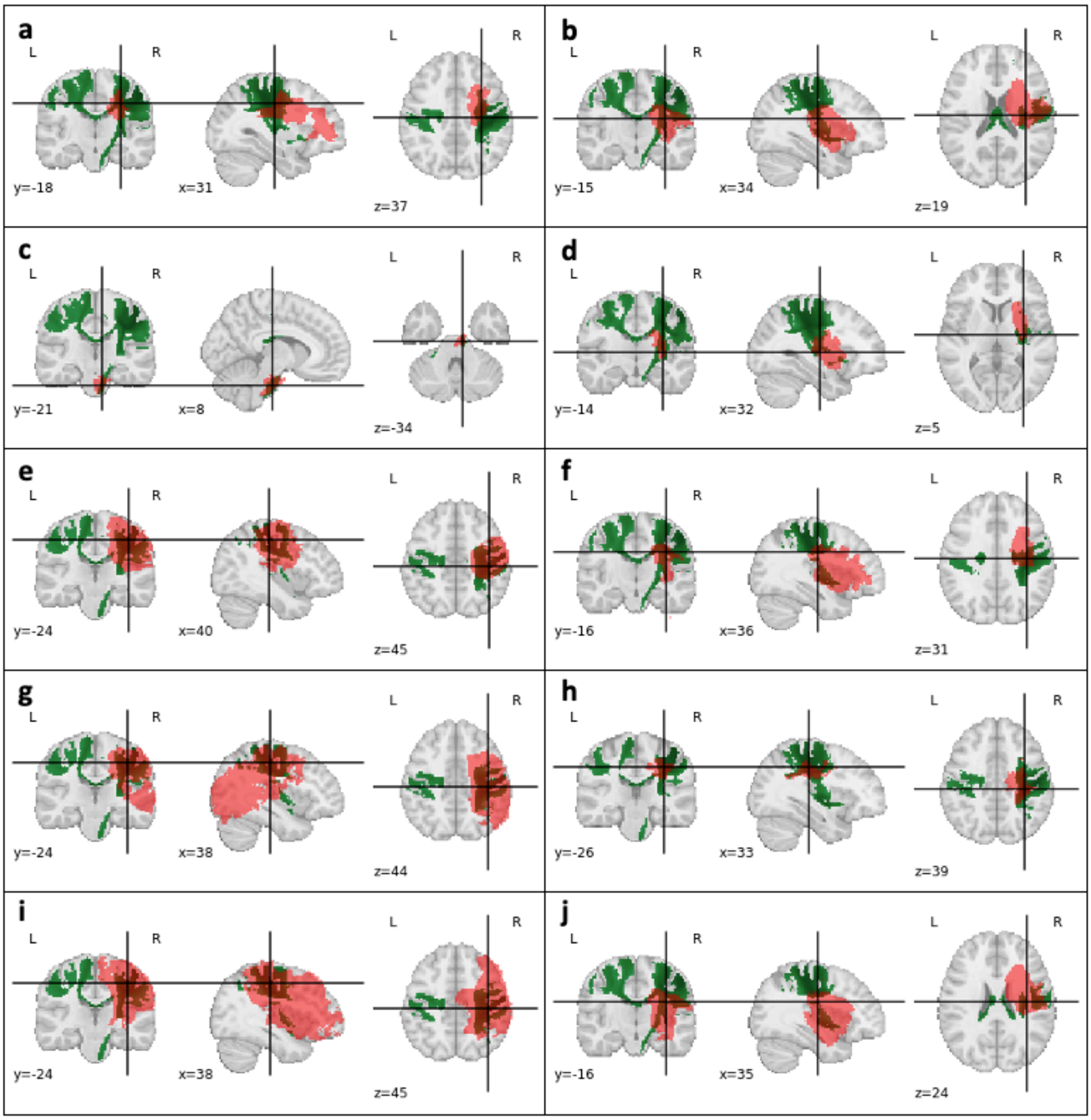

**Supplementary figure 32**: Patients with strong left upper-limb motor control deficit (part 1/2). **a-j**: In red, the stroke lesion; in green, the somato-motor network for the left hand (RSN 09).

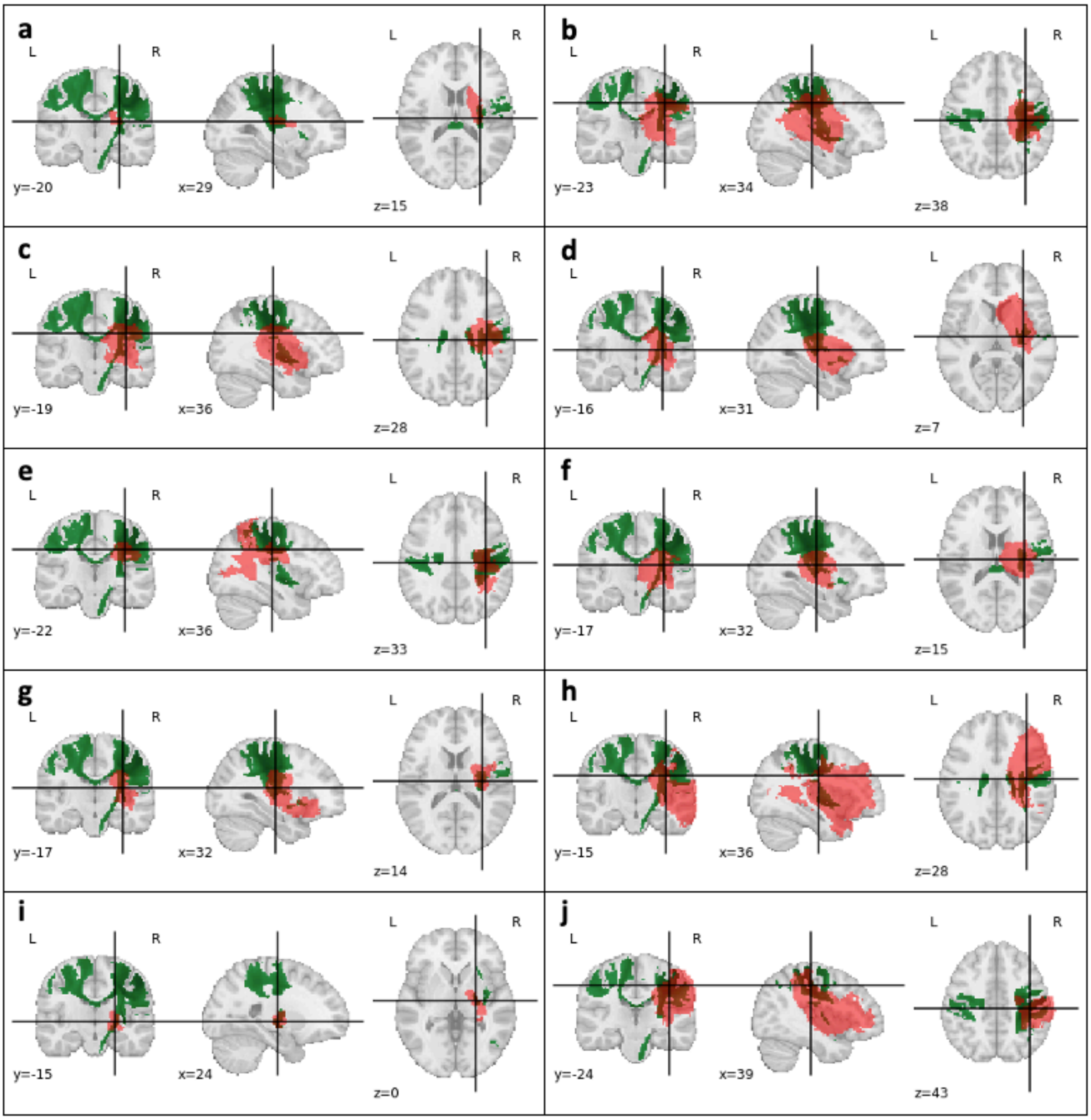

**Supplementary figure 33**: Patients with strong left upper-limb motor control deficit (part 2/2). **a-j**: In red, the stroke lesion; in green, the somato-motor network for the left hand (RSN 09).

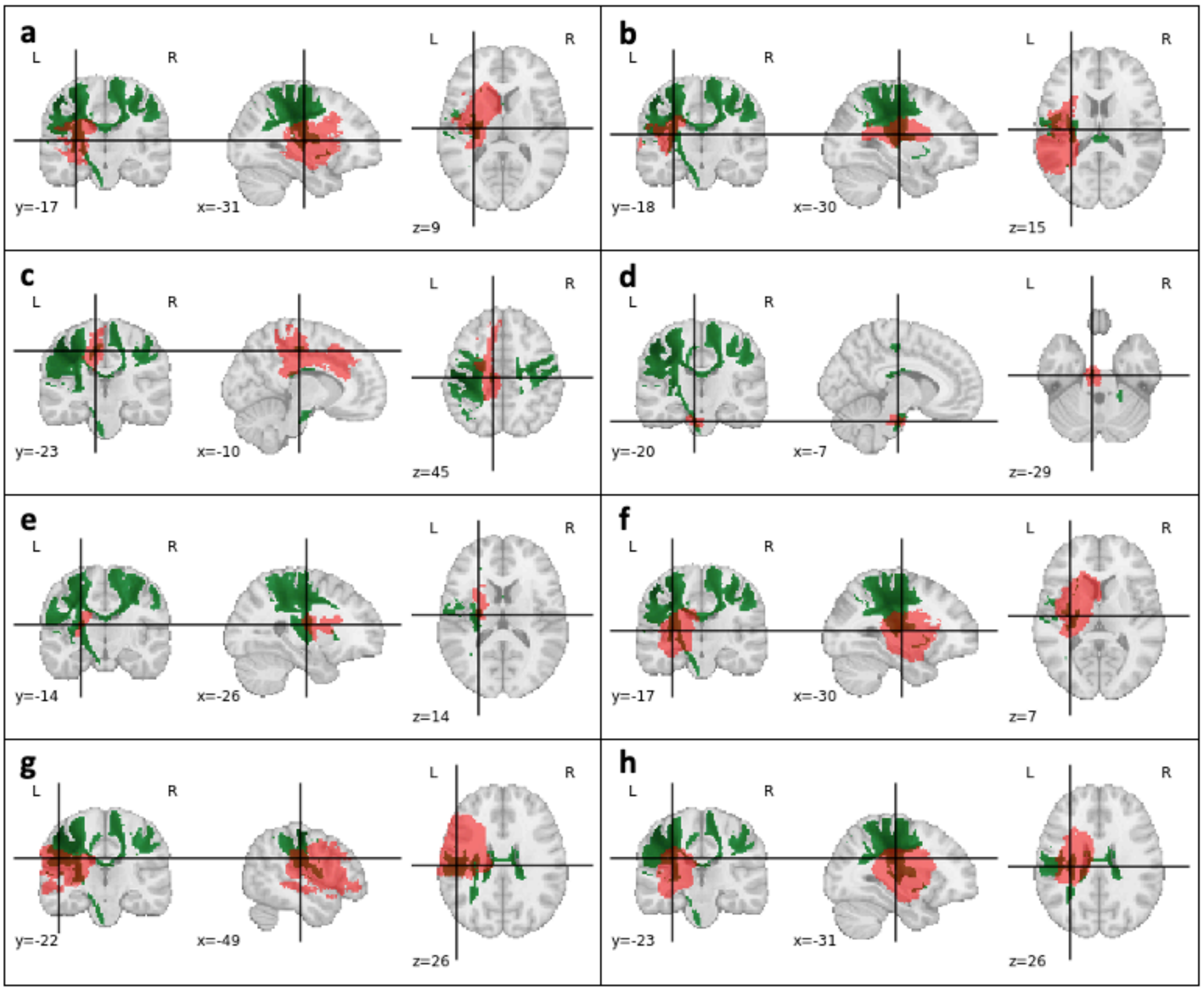

**Supplementary figure 34**: Patients with strong right upper-limb motor control deficit (part 1/2). **a-h:** In red, the stroke lesion; in green, the somato-motor network for the right hand (RSN 08).

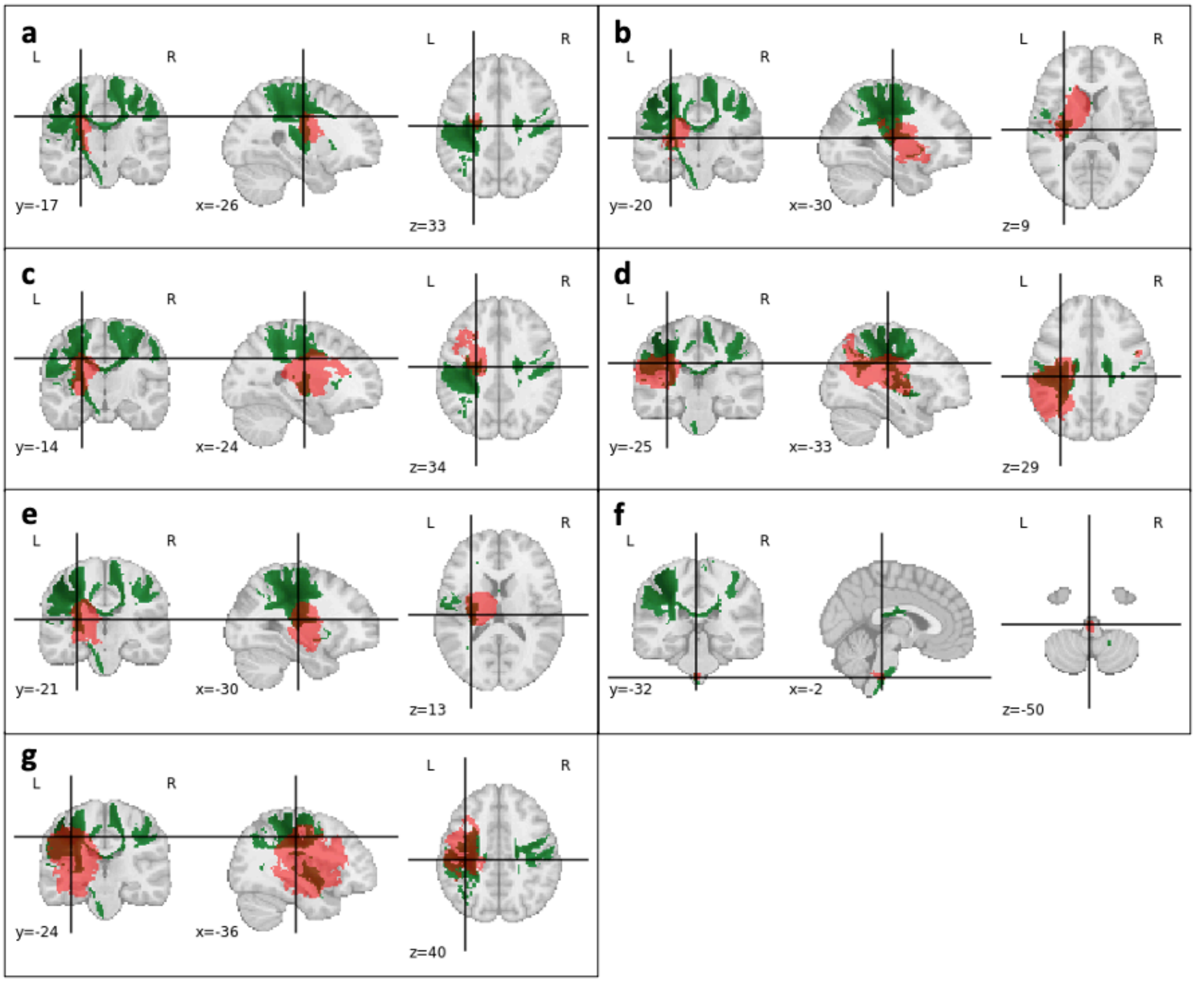

**Supplementary figure 35**: Patients with strong right upper-limb motor control deficit (part 2/2). **a-g:** In red, the stroke lesion; in green, the somato-motor network for the right hand (RSN 08).

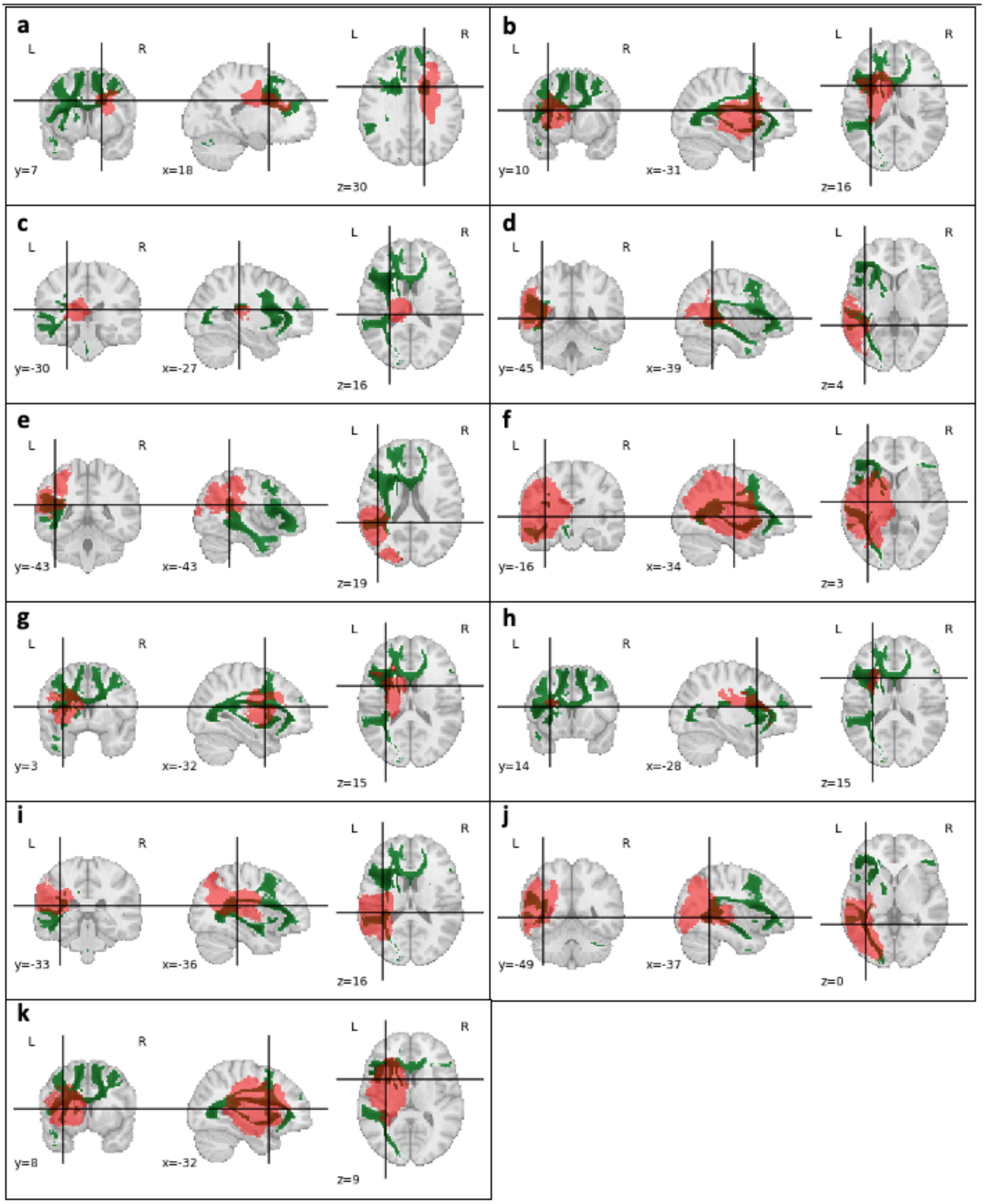

**Supplementary figure 36**: Patients with strong right language deficit (part 1/2). **a-k:** In red, the stroke lesion; in green, the language production network (RSN 20).

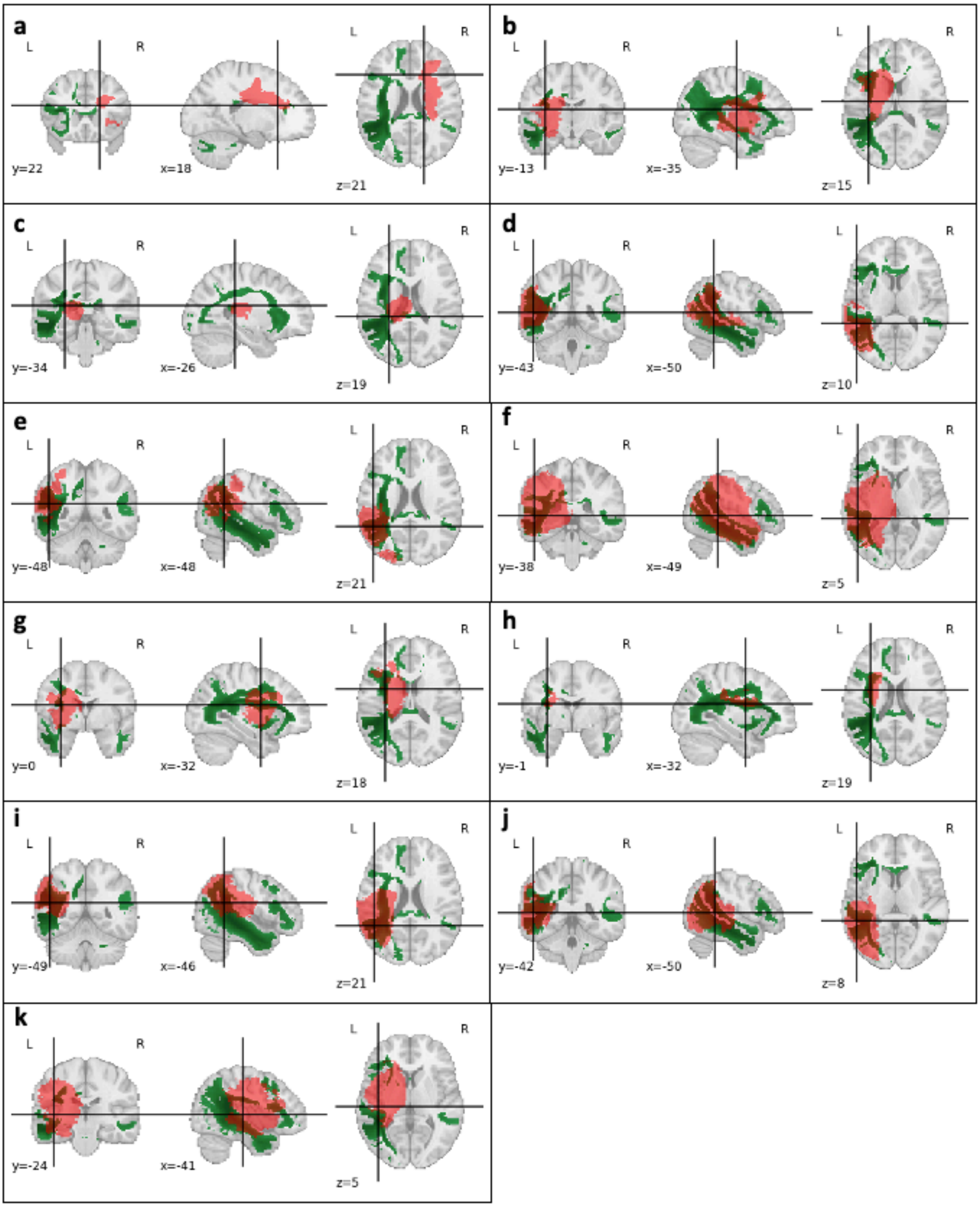

**Supplementary figure 37**: Patients with strong right language deficit (part 2/2). **a-k:** In red, the stroke lesion; in green, the language comprehension network (RSN 25).

#### Overlap of RSNs and plurality of symptoms

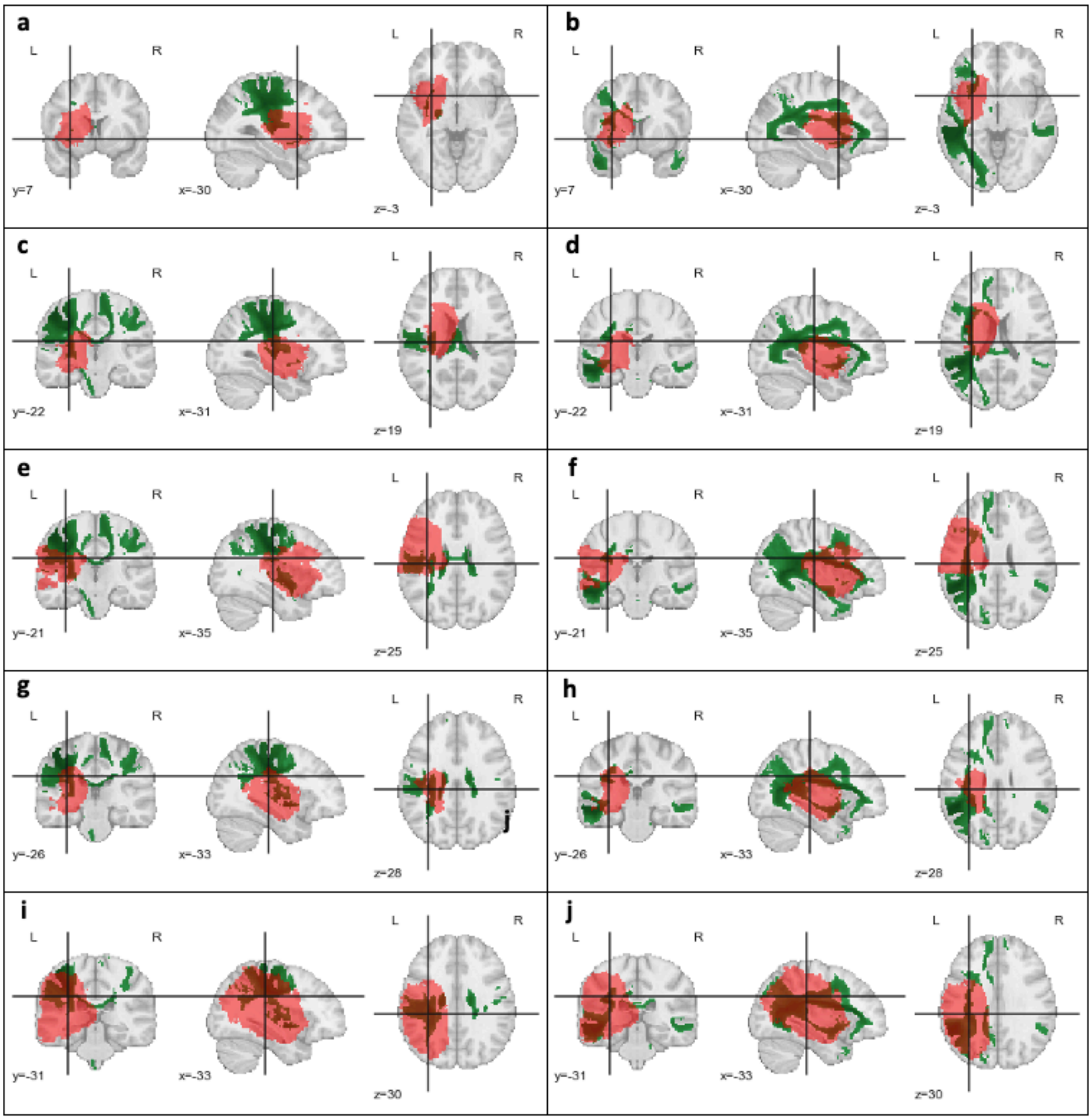

**Supplementary figure 38**: Patients with strong right language deficit (part 1/2). **a-j:** One patient (lesion) per row. In red, the stroke lesion; in green, the right hand somato-motor network (RSN 08) for the left column, and the language comprehension network (RSN25) for the right column.

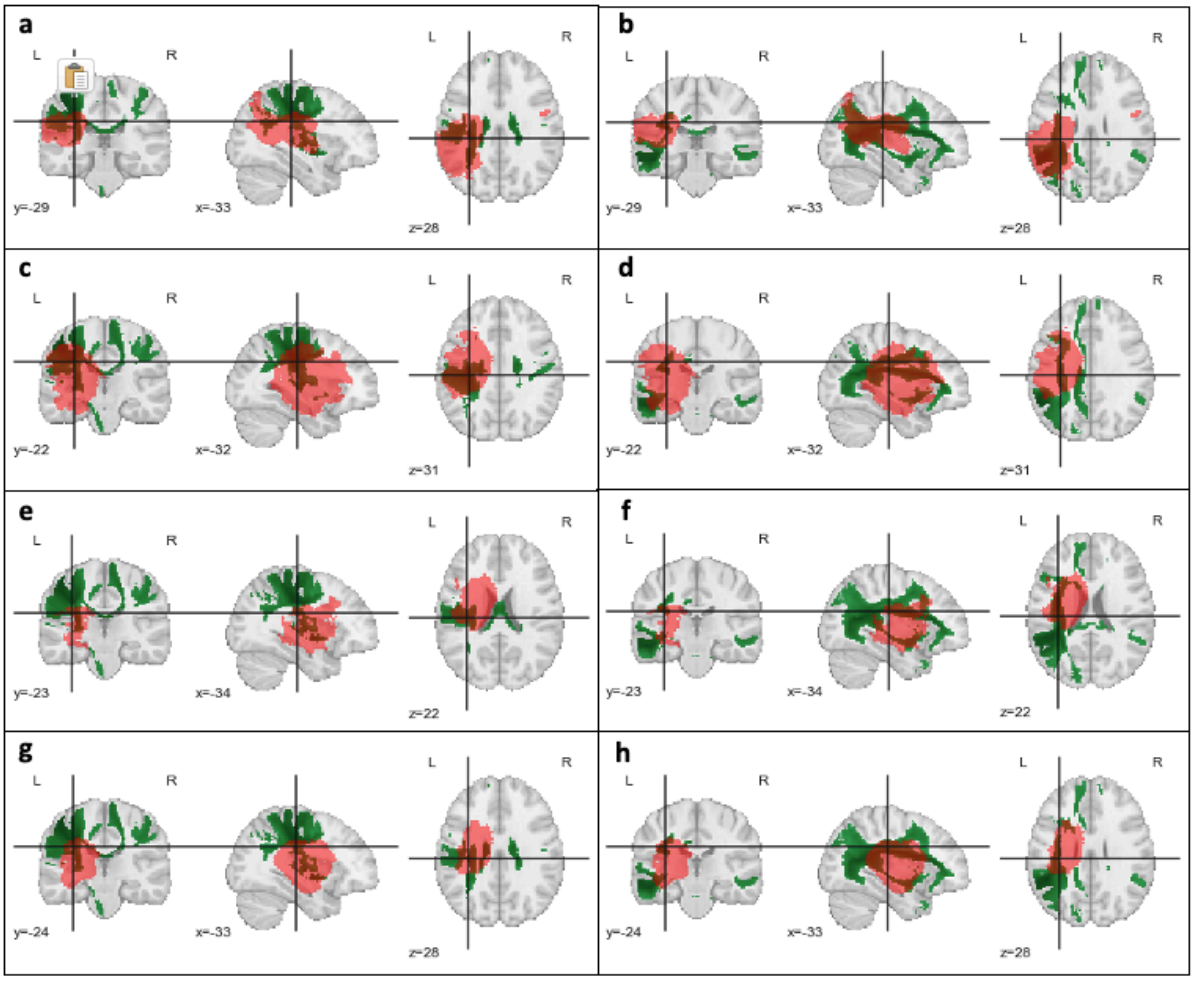

**Supplementary figure 39**: Patients with strong right language deficit (part 2/2). **a-h:** One patient (lesion) per row. In red, the stroke lesion; in green, the right hand somato-motor network (RSN 08) for the left column, and the language comprehension network (RSN25) for the right column.

#### WhiteRest manual

WhiteRest is a module of the Functionnectome software developed to help you explore the WhiteRest atlas and analyze the potential impact of a white matter lesion on RSNs.

*For now, WhiteRest is only available through command line call, but we have plans to create a user interface and a web app to facilitate its use.*

##### Disclaimers

No financial conflicts.

Not licensed for medical use.

The software and its codes are licenced under the GNU General Public License.

##### Installation

- A Python environment is needed to use WhiteRest. For detailed instructions, check the start of our video tutorial for the Functionnectome on YouTube.
- WhiteRest is installed along the Functionnectome, so following the instruction in the Functionnectome manual will install the software.
  The following commands in a terminal with a proper Python environment will install it:
  pip install git+https://github.com/NotaCS/Functionnectome.git
  *or*
  python -m pip install git+https://github.com/NotaCS/Functionnectome.git
- Download and unzip the WhiteRest atlas and RSN labels information following the link: https://www.dropbox.com/s/mo4zs159rqhqopv/WhiteRest.zip?dl=0
  - Note that the white matter atlas file is the same as the one downloadable from NeuroVault (https://neurovault.org/collections/11895/), so if you want to use the grey matter atlas, you can download it from there too (https://neurovault.org/collections/11937/).
- After the installation, to check if WhiteRest was properly installed, type "WhiteRest -h" (without the quotation marks) in the terminal and press Enter. The description and the help should be displayed.

##### Using WhiteRest

- WhiteRest can compute two types of scores: the **DiscROver** score, and the **Presence** score. The **DiscROver** score is the default option, it estimates the disruption of RSNs by a brain lesion, using Disconnectome maps (method available in the BCBtoolkit, www.bcblab.com, see Foulon et al. 2018). The **Presence** score measures locally, in a ROI, which and how much of each RSNs are present. For more detailed about the computation of the scores, refer to the ***Score computation*** section below.
- Prepare the lesion mask (or region of interest) file you wish to use. The file must be in NIfTI format (.nii or .nii.gz), in MNI space, and with 2mm$^3$ isotropic voxels. If you want to compute the DiscROver score, you can also prepare the disconnectome of the lesion, or use the quickDisco algorithm embedded in WhiteRest (see the ***Score computation*** section below) by simply giving it the lesion mask (that's the default option).

- Open a terminal. If necessary, activate the Python environment where the Functionnectome is installed (if it is not the default one).
- Type “WhiteRest” followed by the paths to paths to the files and the chosen options and press Enter.
  The minimal inputs should be (in that order):
  - The path to the ROI file you wish to explore (as a nifti file in the MNI space)
  - The path to the white matter maps of the WhiteRest atlas
  - The path to the RSN labels information from the atlas

  For example:
  “WhiteRest /myhome/my_ROI.nii.gz /myhome/WhiteRestAtlas_WM.nii.gz /myhome/WhiteRest_labels.txt”

By default, WhiteRest will output a table with the computed score for each RSN. If no output file is given (with the "-ot" option), the table will be printed in the terminal. Otherwise, it can be saved as a text file (.txt or .csv), which can be imported to a spreadsheet software (such as Excel) for further processing.

**If you are computing the DiscROver score** and only giving the **lesion mask** (which is the default expected behavior), WhiteRest will need to compute the disconnectome of the lesion. It will do it using the quickDisco algorithms that employs white matter priors from the Functionnectome to estimate the disconnection pattern. For that, you will also need to download these white matter priors too (if it’s not already the case). If you don’t have them, an error message will pop-up explain how to do it, but essentially you need to launch the Functionnectome interface (command line: *FunctionnectomeGUI*), select the priors you want (in the “Choice of priors” drop-down menu), the V1.D.WB being the same as the one used to create the atlas. Then click on “Manual download” and follow the instructions.
Also, because computing a disconnectome can be a bit slow (though much faster with quickDisco than with the traditional method), ranging from a few second to a few minutes depending on the size of the lesion, **you can speed the process up** by parallelizing the task. Simply use the “**--multiproc”** (or just “-m”) option, followed by a number, to specify the number of processes to launch in parallel. You can **save the generated disconnectome** for later use with the “**--out_disco**” (or “-od”) option and giving a file-path to where to save the output file (in Nifti format).
**If you are computing the DiscROver score** and **already have the disconnectome** on hand, you can skip the process and replace the input ROI by the disconnectome. You just need to specify it simply by adding the “**--disco**” (or “-d”) option in command line.
**If you are computing the Presence score**, you will need to specify it in the “--score” (or “-s”) option, by adding either “presence” (if you only want the Presence sore) or “both” (if you want Presence and DiscROver scores) after the option.

All WhiteRest options and arguments:

```
positional arguments:
  in_ROI          Path of the ROI file (.nii, .nii.gz).
  atlas_maps      Path of the RSN atlas (.nii, nii.gz).
  atlas_labels    Path to the atlas labels identifying the RSNs.

optional arguments:

  -h, --help
           show this help message and exit

  -s SCORE, --score SCORE
           The score(s) to be computed. Can be "discrover",
           "presence", or "both" (default is "discrover")

  -ot OUT_TABLE, --out_table OUT_TABLE
           Path to save the score results (.txt or .csv).

  -od OUT_DISCO, --out_disco OUT_DISCO
           Path to save the lesion's disconnectome (.nii or
           .nii.gz), if computed.

  -z Z_THRESH, --Z_thresh Z_THRESH
           Threshold to apply to the atlas z-maps (default z>7).

  -b, --binarize
           Binarize the maps after thresholding.

  -p OUT_PIE, --out_pie OUT_PIE
           Path to save a pie-chart figure of the results (.png).

  -pt THR_LOW_PIE, --thr_low_pie THR_LOW_PIE
           DiscROver % under which the RSNs are grouped on the
           pie-chart (default <5%).

  -d, --disco
           To be specified when the "in_ROI" input given is not
           the lesion but the disconncetome of the lesion.
           Incompatible with Presence score computation.

  -m MULTIPROC, --multiproc MULTIPROC
           Number of processes to run in parallel (default = 1).
```

Example of command lines:

- Computation of DiscROver score from a lesion:

WhiteRest /myhome/myProject/my_lesion.nii.gz /myhome/WhiteRestAtlas_WM.nii.gz /myhome/WhiteRest_labels.txt -ot /myhome/myProject/table_results.txt -m 4 -od /myhome/myProject/my_lesion_disco.nii.gz

- Computation of DiscROver score from a disconnectome (not the "-d" at the end):

WhiteRest /myhome/myProject/my_lesion_disco.nii.gz /myhome/WhiteRestAtlas_WM.nii.gz /myhome/WhiteRest_labels.txt -ot /myhome/myProject/table_results.txt -d

- Computation of the Presence score:

WhiteRest /myhome/myProject/my_ROI.nii.gz /myhome/WhiteRestAtlas_WM.nii.gz /myhome/WhiteRest_labels.txt -s presence -ot /myhome/myProject/table_results.txt

#### Score computation

##### DiscROver score

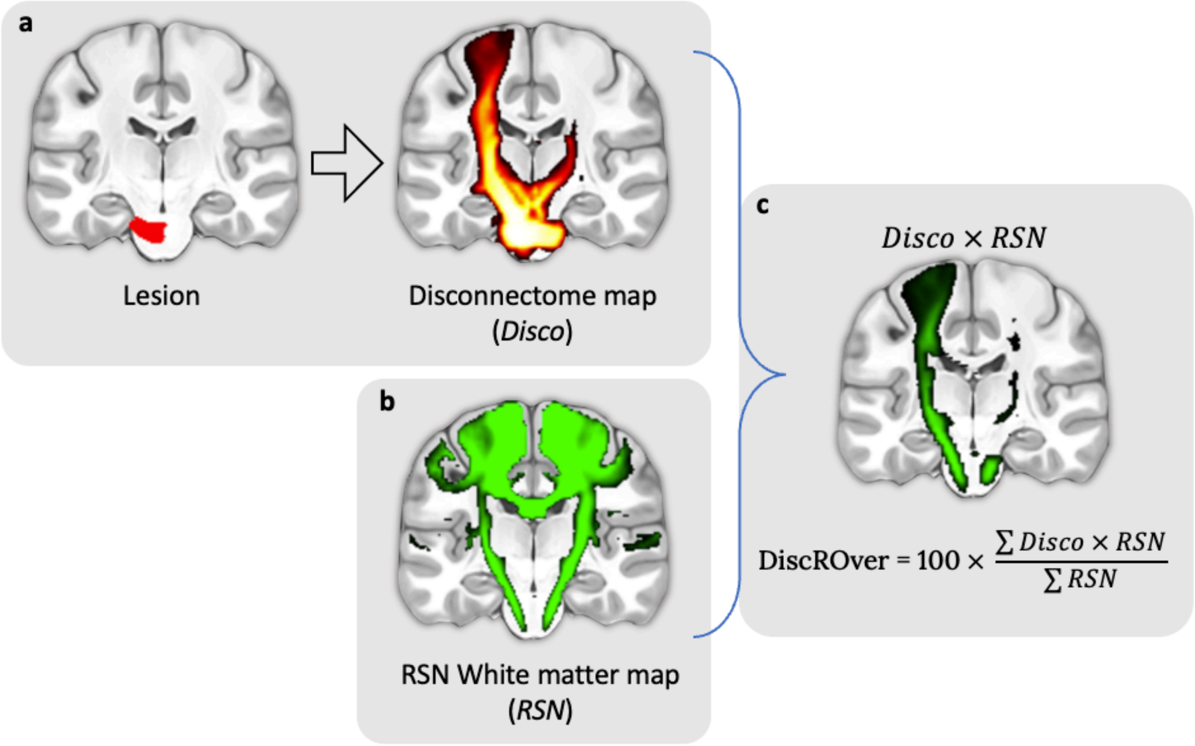

**Figure**: Steps for the computation of the DiscROver score. **a** - Lesion mask (left) and associated disconnectome (right). **b -** RSN map used for the DiscROver score computation. **c** - Visual representation of the weighted overlap, and computation of the DiscROver score. Disco: Disconnectome map; RSN: Resting-state network map.

DiscROver stands for "Disconnectome RSN Overlap". For a given RSN and a given lesion, the DiscROver score is computed as follows: First, the extent of white matter fibres disconnected by the lesion is estimated using the Disconnectome method. This method yields a disconnectome map displaying the probability of structural connectivity between the lesion and each voxel of the brain (Fig. a). Hence, the higher the value on the disconnectome map,

the more likely the disruption of connectivity in the voxel due to the lesion. Then, the weighted overlap of the RSN (Fig. b) with the disconnectome is computed by voxel-wise multiplication of the RSN map and the disconnectome map (Fig. c). The DiscROver score is computed as the sum of the values of this weighted overlap map, normalised by the sum of the values in the RSN, and multiplied by 100. With this score, 0 means that the lesion does not impact any white matter voxel part of the RSN, 100 means that it impacts the whole RSN.
The complete computation of the DiscROver score is summarised here:

$$DiscROver(RSN, Disco) = 100 * \frac{\sum_{v \in RSN} Z_{RSN}(v) \times P_{Disco}(v)}{\sum_{v \in RSN} P_{Disco}(v)}$$

With "*RSN*" the atlas white matter Z-map of a given RSN, with its voxel values annotated as "$Z_{RSN}(v)$", and "*Disco*" the disconnectome map of a given lesion, with its voxel values annotated as "$P_{Disco}(v)$".

**In the output** from WhiteRest, the DiscROver score is in the column "DiscROver (%)", while the "DiscROver (raw)" column results from the non-normalised score computed with $\sum_{v \in RSN} Z_{RSN}(v) \times P_{Disco}(v)$ in the above equation.

**QuickDisco**
To compute ease the computation of the disconnectome required to compute the DiscRover score, we introduced the **quickDisco algorithm**. It is used by the WhiteRest program but can also be used independently, as a module of the Functionnectome.
Simply type "**quickDisco -h**" in the terminal to display the help text for this program. In short, where the original Disconnectome method uses tractography to estimate the connectivity between a brain lesion and the rest of the brain (thus revealing the disconnection pattern of the lesion), quickDisco uses the white matter priors (derived from 100 tractograms) to do the same. The priors contain the connectivity map from every voxel of the brain, so using them to compute the disconnectome is relatively straightforward: it is the maximum projection, across the map of all the voxels in the lesion, for each brain voxel. For a given lesion, the equation would be:

$$P_{Disco}(v) = Max_{l \in lesion}(Pmap_l(v))$$

With $v$ a given brain voxel, $P_{Disco}$ the disconnectome (giving the probability of disconnection at each voxel), $lesion$ the lesion in question, $l$ one voxel of the lesion, and $Pmap_l(v)$ the probability of connectivity between the voxel $l$ and the voxel $v$, as per the priors (or, if you prefer, the value of the connectivity map of $l$ ($Pmap_l$), at the voxel $v$).

#### Presence Score

The Presence score measures the involvement, or "proportional presence" ($Presence_{prop}$) of each RSN in a ROI by first computing a "raw presence" ($Presence_{raw}$), adding up the z-score value of all the RSN white matter map voxels in the ROI (Equation 1). This raw presence score is then normalised by dividing it by the sum of all z-scores in the RSN

(Equation 2), and finally converted into a proportional presence by dividing it by the sum of the normalised presence score of all the RSNs sharing part of the ROI (Equation 3):

For a given RSN and a given ROI,

$$Presence_{raw}(RSN, ROI) = \sum_{v \in ROI} z_{RSN}(v) \quad [1]$$

$$Presence_{norm}(RSN, ROI) = \frac{Presence_{raw}(RSN,ROI)}{\sum_{v \in RSN} z_{RSN}(v)} \quad [2]$$

$$Presence_{prop}(RSN, ROI) = \frac{Presence_{norm}(RSN,ROI)}{\sum_{rsn \in Atlas} Presence_{norm}(rsn,ROI)} [3]$$

with $v$ a voxel from the ROI, $z_{RSN}(v)$ the z-score of the voxel $v$ in the white matter map of the studied RSN.

All these metrics are proposed in the output table for a more complete analysis of the atlas:

- The RSN normalised presence ***Presence/RSN (%)***: The fraction of presence in the ROI of an RSN compared to the whole white matter map of the said RSN (i.e. how much of the RSN is in the ROI). (Eq. 2)
- The proportional presence ***Presence prop. (%)***: Proportion of the presence of one RSN compared to the sum of the presence of all RSN in the ROI (Eq. 3)
- The “raw” presence score ***Presence (raw)***: Sum of the z-score of all the voxels from the white matter map of one RSN in the ROI (Eq. 1)
- The ROI coverage ***Coverage (%)***: The volume fraction of the ROI occupied by the RSN.

Essentially, the normalised presence score reflects how much of a network is intersecting with the ROI, and the proportional presence score can be used to compare the involvement of the affected RSNs. The Coverage indicates how much of the ROI is covered by a RSN.
